# Lineage-specific genes are clustered with allorecognition loci and respond to G × E factors regulating the switch from asexual to sexual reproduction in *Neurospora*

**DOI:** 10.1101/2022.06.10.495464

**Authors:** Zheng Wang, Yaning Wang, Takao Kasuga, Yen-Wen Wang, Francesc Lopez-Giraldez, Yang Zhang, Zhang Zhang, Caihong Dong, Anita Sil, Frances Trail, Oded Yarden, Jeffrey P. Townsend

**Affiliations:** Department of Biostatistics, Yale School of Public Health, New Haven, Connecticut 06511, USA; Institute of Microbiology, Chinese Academy of Sciences, Beijing 100101, China; College of Biological Sciences, University of California, Davis, CA95616, USA; Yale Center for Genomic Analysis, New Haven, Connecticut 06511, USA; National Genomics Data Center, Beijing Institute of Genomics, Chinese Academy of Sciences, Beijing 100101, China; Department of Microbiology and Immunology, University of California, San Francisco, CA94115, USA; Department of Plant, Soil and Microbial Sciences, Michigan State University, East Lansing, MI 48824, USA; Department of Plant Pathology and Microbiology, The Robert H. Smith Faculty of Agriculture, Food and Environment, The Hebrew University of Jerusalem, Rehovot, Israel; Department of Ecology and Evolutionary Biology, Program in Microbiology, and Program in Computational Biology and Bioinformatics, Yale University, New Haven, Connecticut 06511, USA

## Abstract

Lineage-specific genes (LSGs) have long been postulated to play roles in the establishment of genetic barriers to intercrossing and speciation. However, there is a lack of working hypotheses as to how they might play that role. In the genome of *Neurospora crassa*, most of the 670 *Neurospora* LSGs that are aggregated adjacent to the telomeres are clustered with 61% of the HET-domain genes, which regulate self-recognition and define vegetative incompatibility groups. Among the 342 LSGs that are dynamically expressed during both asexual and sexual phases, 64% were detectable on unusual carbon sources such as furfural and HMF—wildfire-produced chemicals that are a strong inducer of sexual development. Expression of a significant portion of the LSGs was sensitive to light and temperature, factors that regulate the switch from asexual to sexual reproduction. Furthermore, expression of the LSGs was significantly affected in the knockouts of *adv-1* and *pp-1* that regulate hyphal communication, and expression of more than one quarter of the LSGs was affected by perturbation of the mating locus. Accordingly, we propose a gene-by-environment interaction model encouraging further investigation of the roles of LSGs and HET-domain genes in speciation in *Neurospora*. This gene-by-environment interaction model emphasizes the roles of the LSGs in response to genetic and environmental factors, leading to the regulation of the switch from the asexual growth and fusion, such that vegetative incompatibility governed by allorecognition promotes allelic homogeneity, sexual reproduction, and outbreeding, whereas VI repression and meiotic recombination promotes allelic polymorphism.

## Introduction

Since the emergence of life, molecular evolution has contributed to the accumulation of novel and diverse features in all kinds of organisms. Two fundamental components of that molecular evolutionary novelty are new genes and novel gene functions, which have long been considered to be emergent properties of gene duplication and rearrangement. Nevertheless, genomes often harbor numerous orphan genes or lineage specific genes (LSGs)—novel genes that have no homologues in distantly- or closely-related lineages and that cannot be tracked to ancestral lineages. These LSGs, including de novo genes that evolve from previous non-coding DNA and non-genic elements [1], manifest in large numbers across a diversity of organisms, such that they represent nearly one-third of the genes in all genomes, including phages, archaea, bacteria, and eukaryotic organisms [2]. Three key challenges have thus been the focus of studies of LSGs: how to identify LSGs, how to track their evolutionary histories, and perhaps most importantly, how these genes are integrated into pre-existing gene interaction networks [3].

Accurate identification of LSGs can, at times, be difficult. They originate via three evolutionary processes: (1) rapid gene evolution, which is refractory to tracking homology on the basis of sequence conservation; (2) intragenomic gene loss and gain or horizontal gene transfer, which can convey higher fitness in response to genetic/environmental changes; and (3) accumulated mutations that establish novel function, evolving slowly but long enough on an independent lineage that the gene phylogeny cannot be tracked back to its distant ancestor or to related lineages [4–6].

Studies of the genomic characteristics of LSGs in several model organisms have revealed the likely origins of LSGs from gene duplication, non-coding sequences, as well as fast-evolution at conserved genomic positions. One example supporting the frequency of origin by rapid divergence after gene duplication and rearrangement can be found in yeast, where the presence of 55% to 73% percent of the LSGs can be explained by sufficient divergence from sister species [6]. Some lineage-specific protein-coding genes might have directly evolved from non-coding regions in the genome [7], as has been described in the tests of fruit flies [8,9]. One hundred seventy-five *de novo* genes in Asian rice corresponded to recognizable non-genic sequences in closely related species [10]. These investigations using model organisms confirmed unique characteristics of these *de novo* genes, making good frameworks for investigating LSGs in other species. However, linkages between these revealed genomic characteristics and the integrative functions of LSGs remain unclear. Therefore, systematic approaches that combine study of comparative genomics with functional assays of gene expression and gene-perturbation phenotypes using well-established model systems are critical to integrate the investigation of the LSGs’ origination and function.

The most frequently used approach to identify LSGs is using phylostratigraphy [11,12]. Precise identification of the origins of *de novo* genes using a phylostratigraphic approach is critically dependent on accurate gene annotation and extensive comparison among proper representative genomes [13]. It is difficult to distinguish whether genes with no homologues in closely related lineages are true LSGs as opposed to lacking homologues in closely related lineages that have few genomes available. An alternative to phylostratigraphy is gene synteny, which compares each gene’s position relative to its neighbors. A recent study suggested that if the neighbors of a gene are in a conserved order in other species, then the gene is likely to correspond to whatever is at the orthologous position in the other species as well—even if the sequences do not match [14].

With detailed pangenomic analyses, identification of LSGs, including *de novo* protein-coding genes, need to be further verified with a systematic approach focusing on possible functional novelty and genetic signals that may be associated with such a novelty [3]. Systematic assessment of the putative LSG function can track their behaviors during the growth and development and verify their possible roles by examination of corresponding knockdown or knockout phenotypes [3]. LSGs are naturally thought to be important to species- or genus-level adaptations of development to taxon-specific ecology.

The well-annotated model species in the genus *Neurospora*, *N. crassa, N. tetrasperma* and *N. discreta* of the class Sordariomycetes, provide a set of three closely-related genomes enabling investigation of possible genetic novelties associated with recent and rapid ecological divergences [15], such as responses to nutrients and other environmental factors and developmental divergences such as heterothallic and pseudohomothallic outcrossing during sexual reproduction as well as incompatibility during vegetative growth. *Neurospora* species are highly adapted to the postfire environment, capable of fast asexual growth and reproduction on simple nutrients and have long been genetic models for eukaryotic metabolic regulation and for mating, meiosis and morphological development during reproduction [16–18]. In fact, fungi in the Sordariomycetes exhibit remarkable ecological, biological, and morphological diversity [19–21]. These fungi also exhibit diverse reproductive modes, including heterothallism, pseudohomothallism, and homothallism with or without asexual reproduction [18,22–26]. Such diversities in ecology and development that have evolved in parallel or convergently within closely related species in a single fungal class, may provide avenues toward an understanding of the G × E (gene-environment interaction) impacts of lineage-specific elements on the evolutionary process. Due to the repeat-induced point mutation (RIP) genome-defense system, *Neurospora* is known to lack recent gene duplications—a major source of evolutionary novelty [27,28]. Nevertheless, comparing representative genomes in prokaryotes, plants & animals, Basidiomycota, major lineages of Ascomycota, and *Chaetomium globosum,* which is closely related to *N. crassa*, 2219 orphan genes were identified in *N. crassa* by phylostratigraphy [29]. In the past decade, many more fungal genomes have been sequenced. Their sequence, along with the advances in genome sequencing and annotation techniques [30–32], provide a more inclusive comparison for identifying lineage specific genes. Therefore, to understand how important the roles that LSGs play in genome-wide regulation during the whole life history, we investigated signals of recent selection pressures in LSGs, focusing on genes involved in environmental responses and fertility or mating in *Neurospora*.

We also observed that several LSGs are allorecognition loci, often referred to as *het* (heterokaryon incompatibility) or *vic* (vegetative incompatibility) genes. As their names indicate, *het* or *vic* genes regulate allorecognition during vegetative growth, and only individuals with compatibility at all of their *het*-loci can fuse and simply expand their colonies [33]. Some HET-domain genes have pleiotropic effects in sexual development in some fungal species and play direct roles in reproductive isolation and speciation within sympatry [34]. HET-domain genes can be predicted with conserved HET domain in protein sequences, and there were 69 HET-domain genes reported (of which 68 were mapped to the original genome-sequenced strain of *N. crassa*) [33]. Chromosomal locations that are itinerant over evolutionary time, high duplication rates and high frequencies of gain and loss are shared traits of HET-domain genes and LSGs, suggesting the merits of integrative investigations of both gene groups.

## Results

### Summary

In this study, we identified and verified 670 lineage-specific genes (LSGs) in *Neurospora crassa* via BLAST search against inclusive representative genomes using previously published phylostratigraphy data that was based on a limited set of representative genomes. Using two clustering approaches, we discovered that over 60% of the 670 LSGs formed clusters in the telomere regions and clustered with the HET-domain genes. However, most of the LSGs are not functionally annotated (e.g., with gene ontology terms). Therefore, to assess the possible functional roles of the LSGs, we analyzed genome-wide gene expression data on *N. crassa* growing on distinct media at different stages of life history and under different light and temperature conditions. We observed that nearly half of the LSGs were actively expressed during asexual and sexual growth. A substantial number (291) LSGs were induced by the presence of furfural. 158 LSGs were exclusively expressed in furfural cultures, in contrast to only 17 LSGs that were exclusively expressed in cultures on media supplied with common simple carbohydrates. We also reported a significant portion of the LSGs being turned on or off by the changes of light exposure and temperature, conditions that are critical for *N. crassa* asexual and sexual reproduction. We further examined expression of the LSGs and HET-domain genes in knockouts of two transcription factors, *adv-1* and *pp-1*. Both transcription factors play multiple roles in asexual and sexual development in *N. crassa*. In addition, we sequenced and analyzed genome-wide gene expression in a loss-function mutant at the mating locus *mat 1-2-1* in a *mat a* strain during the crossing. The LSGs were more likely to be affected by gene-manipulation than other genes, compared with other non-LSG genes and HET-domain genes. They were more likely to be turned on or turned off completely, rather than being turned up or down slightly. Finally, we examined asexual and sexual growth phenotypes for 367 available KO strains of the *Neurospora* LSGs. We identified two LSGs with abortive sexual reproduction, and several LSGs with minor phenotypes in response to high temperature or mycelium morphology.

### 670 LSGs were identified in Neurospora genomes

We defined *Neurospora* lineage-specific genes (LSGs) as *N. crassa* genes exhibiting homology to only to genes found in sequences from species within the genus *Neurospora* [11]. To identify *Neurospora* LSGs, phylostratigraphy was performed on representative taxa for major fungal lineages and several non-fungal reference genomes, and 1872 *N. crassa* genes were identified as putative LSGs (**Fig. 1, Table S1**). Within these 1872 *N. crassa* genes, a total of 695 genes are shared between the genomes of *Neurospora* and the sole species *S. macrospora* in the sister genus **(Table S1**). Further reciprocal-BLAST searches were made for the 1872 genes against available Sordariomycetes genomes, including genomes of *Neurospora* closely related species in *Podospora*, *Pyricularia* and *Ophiostoma* as well as species within the genus including *N. tetrasperma* and *N. discreta* at the National Center for Biotechnology (NCBI) and FungiDB genome database. We identified 670 genes that are *N. crassa* lineage-specific genes (LSGs) (**Fig. S1**, Table S1). There are 7400 single-copy orthologs shared among the genomes of *N. crassa*, *N. discreta,* and *N. tetrasperma* (**Fig. S1A**). Among the 670 LSGs identified in the *N*. *crassa* genome, 241 are unique to *N. crassa* and 405 are shared between *N. crassa* and *N. tetrasperma* with 248 showing no orthologs in *N. discreta*. 181 *Neurospora* LSGs are shared between *N. crassa* and *N. discreta* with 26 showing no orthologs in *N. tetrasperma* (**Fig. S1B**). Because of the special status of *N. crassa* as a model species, here we specifically investigate and report as LSGs those that are specific to *N. crassa* in this study. With more genomes in this fungal class being sequenced and annotated and more non-classified genes in the phylostratigraphy being analyzed, these numbers are expected to be slightly changed. Of the 670 *Neurospora* LSGs, 515 have at least one intron (average: ∼2 introns, maximum: 8 introns in NCU07480). These LSGs encoded proteins ranging from 26 to 1310 amino acids (NCU05561 and NCU04852, respectively). The average LSG length (∼192 amino acids) was significantly shorter than the average length of the non-*Neurospora* lineage-specific genes (non-LSG genes, ∼528 amino acids).

**Figure 1.**
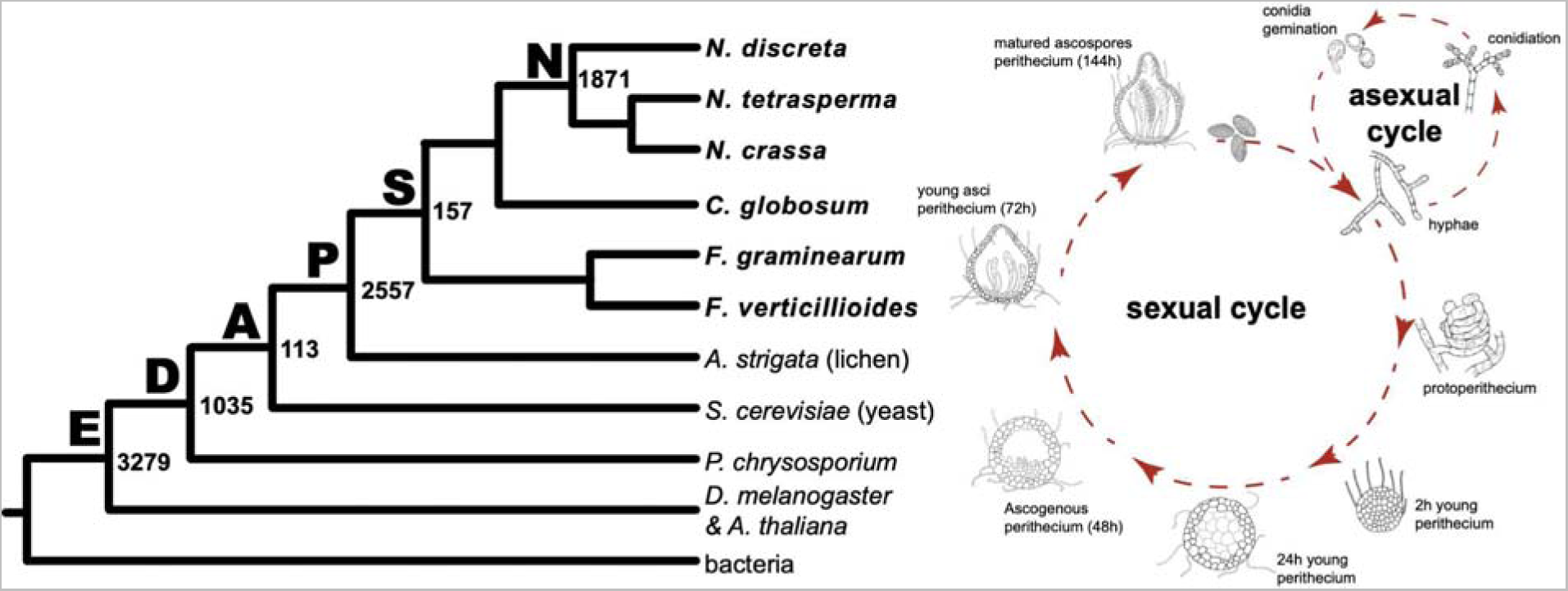
Systematics of *Neurospora* lineage-specific genes (LSGs) and potential roles in fungal growth and development. (A) Genomic phylostratigraphy of lineage-specificity classifications of predicted protein-coding genes (enumerated at ancestral nodes: 3279 Eukaryote-core, 1035 Dikarya-core, 113 Ascomycota-core, 2557 Pezizomycotina-specific, 157 Sordariomycetes-specific, and 1871 Neurospora-specific) that are present within the *N. crassa* genome; (B) Life history of *N. crassa*. These developmental processes have been transcriptionally profiled [dashed red arrows; 35].

**Table 1.** LSGs and HET-domain genes are present in clusters in the *N. crassa* genome.

| Cluster size | Lineage-specific genes (LSGs)<sup>a</sup> |  |  | LSG and HET-domain genes<sup>b</sup> |  |  |
| --- | --- | --- | --- | --- | --- | --- |
|  | # Clusters | # Genes | % Genes | # Clusters | # Genes | % Genes |
| 2 | 57 | 114 | 17.1% | 63 (15 *het*) | 126 | 17.1% |
| 3 | 28 | 84 | 12.6% | 28 (7 *het*) | 84 | 11.4% |
| 4 | 8 | 32 | 4.8% | 12 (6 *het*) | 48 | 6.5% |
| 5 | 14 | 70 | 10.5% | 13 (3 *het*) | 65 | 8.8% |
| 6 | 5 | 30 | 4.5% | 11 (7 *het*) | 66 | 8.9% |
| 7 | 3 | 21 | 3.2% | 2 | 14 | 1.9% |
| 8 | 2 | 16 | 2.4% | 2 | 16 | 2.2% |
| 9 | 3 | 27 | 4.1% | 1 | 9 | 1.2% |
| 10–11 | 1 | 11 | 1.7% | 2(2 *het*) | 21 | 2.9% |
| 16–18 | 1 | 16 | 2.4% | 2(2 *het*) | 35 | 4.8% |
| 23 | 1 | 23 | 3.5% | 1 | 23 | 3.1% |
| Total | 123 | 444 | 66.8% | 129 (42 *het*) | 474 (42 *het*) | 64.4% |

<sup>a</sup> 670 genes; <sup>b</sup> 736 genes (3 *HET-domain* genes are LSGs).

### LSGs are aggregated in the telomere regions and clustered with the HET-domain genes

*Neurospora* LSGs are distributed in all seven chromosomes of the genome, with paralogs from duplicates often clustered together (**Fig. 2**). Window-free maximum-likelihood model averaging of the gene-regionalized probability of *Neurospora* LSGs and HET-domain genes revealed that LSGs were clustered, with significant (*P* < 0.05) clustering on chromosomes I, II, III, IV, and V. Large LSGs clusters are typically aggregated toward the telomeres of each chromosome and frequently contain large non-coding spaces, especially in chromosomes I, III, IV, V, VI, and VII (**Fig. 2 A–G, Table S2**). Detailed clustering revealed with the Cluster Locator [36] tallied 67% of LSGs as present in clusters with a max-gap of five (48% with a max-gap of one, i.e., separated by one gene). About 30% of LSGs were in clusters with more than four genes (**Tables 1 & S3**), including six clusters hosting 9, 10, 16, and 23 genes.

**Figure 2.**
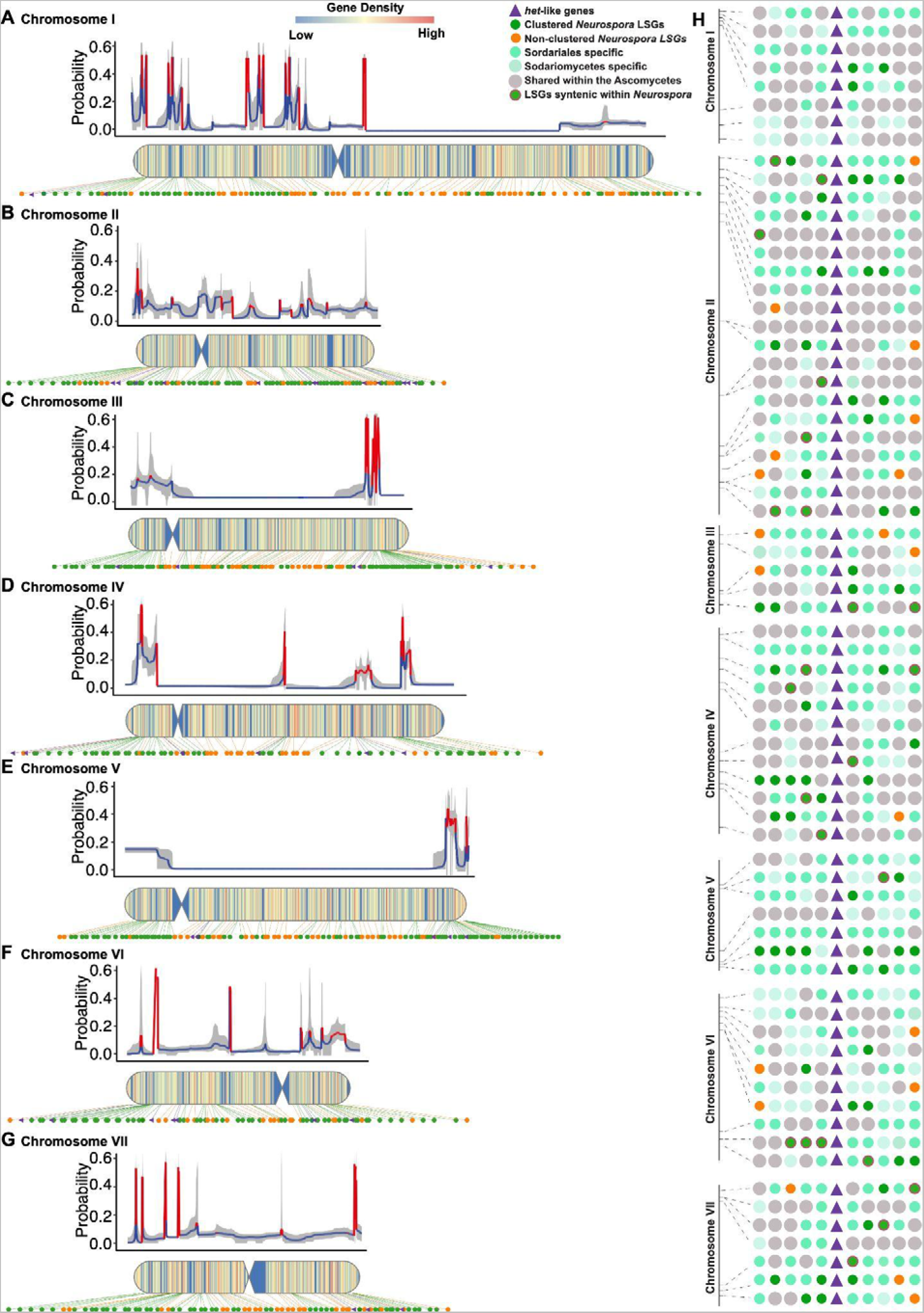
Identification of LSG and LSG-Het clusters in the *N. crassa* genome. Regionalized probability of the lineage-specific status of a gene inferred using MACML, a window-free maximum-likelihood model-averaging approach, heat maps of clustering, and dot-plot of LSGs distributed across (A) chromosome I, (B) chromosome II, (C) chromosome III, (D) chromosome IV, (E) chromosome V, (F) chromosome VI, and (G) chromosome VII. Model-averaged profiles (blue: low clustering of LSGs vs non-LSGs; red: high clustering of LSGs vs non-LSGs; gray shading: 95% model uncertainty interval) quantify the window-free regionalized probability that a gene is an LSG. The heat maps quantify LSG-density across chromosomal windows of 10,000 base pairs. Dots located in accompanying chromosomal heat maps correspond to LSGs (clustered: green [*P* < 0.01], and non-clustered [*P* ≥ 0.01]: orange) and clustered HET-domain genes (purple triangles). (H) Lineage-specificity at four taxonomic depths (greener color intensity corresponds to lineage-specificity at successively lesser taxonomic depths: lineage specificity in *Neurospora* > Sordariales > Sordariomycetes > Ascomycetes; thin orange circle margin indicates synteny within *Neurospora*) of five neighbor genes on either side of 69 HET-domain genes (Table S5) reported in the original genome-sequenced N. crassa strain [[FGSC2489; 38].

**Table 2.**
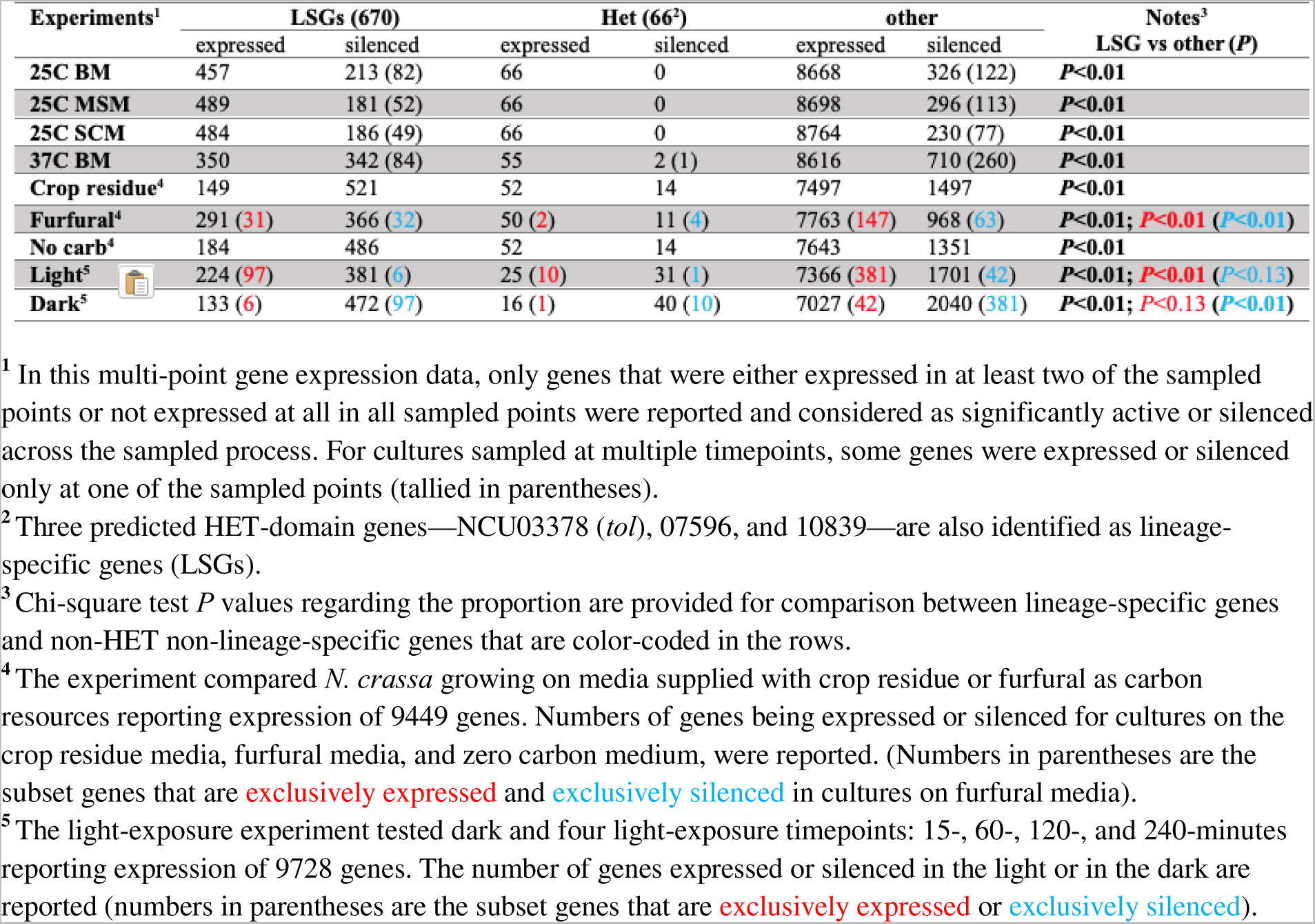
Transcriptomics analysis results on *N. crassa* responses to environmental factors.

| Experiments<sup>1</sup> | LSGs (670) |  | Het (66<sup>2</sup>) |  | other |  | Notes<sup>3</sup><br>LSG vs other (*P*) |
| --- | --- | --- | --- | --- | --- | --- | --- |
|  | expressed | silenced | expressed | silenced | expressed | silenced |  |
| **25C BM** | 457 | 213 (82) | 66 | 0 | 8668 | 326 (122) | *P*<0.01 |
| **25C MSM** | 489 | 181 (52) | 66 | 0 | 8698 | 296 (113) | *P*<0.01 |
| **25C SCM** | 484 | 186 (49) | 66 | 0 | 8764 | 230 (77) | *P*<0.01 |
| **37C BM** | 350 | 342 (84) | 55 | 2 (1) | 8616 | 710 (260) | *P*<0.01 |
| **Crop residue<sup>4</sup>** | 149 | 521 | 52 | 14 | 7497 | 1497 | *P*<0.01 |
| **Furfural<sup>4</sup>** | 291 (31) | 366 (32) | 50 (2) | 11 (4) | 7763 (147) | 968 (63) | *P*<0.01; *P*<0.01 (*P*<0.01) |
| **No carb<sup>4</sup>** | 184 | 486 | 52 | 14 | 7643 | 1351 | *P*<0.01 |
| **Light<sup>5</sup>** | 224 (97) | 381 (6) | 25 (10) | 31 (1) | 7366 (381) | 1701 (42) | *P*<0.01; *P*<0.01 (*P*<0.13) |
| **Dark<sup>5</sup>** | 133 (6) | 472 (97) | 16 (1) | 40 (10) | 7027 (42) | 2040 (381) | *P*<0.01; *P*<0.13 (*P*<0.01) |

<sup>1</sup> In this multi-point gene expression data, only genes that were either expressed in at least two of the sampled points or not expressed at all in all sampled points were reported and considered as significantly active or silenced across the sampled process. For cultures sampled at multiple timepoints, some genes were expressed or silenced only at one of the sampled points (tallied in parentheses).
<sup>2</sup> Three predicted HET-domain genes—NCU03378 (*tol*), 07596, and 10839—are also identified as lineage-specific genes (LSGs).
<sup>3</sup> Chi-square test *P* values regarding the proportion are provided for comparison between lineage-specific genes and non-HET non-lineage-specific genes that are color-coded in the rows.
<sup>4</sup> The experiment compared *N. crassa* growing on media supplied with crop residue or furfural as carbon resources reporting expression of 9449 genes. Numbers of genes being expressed or silenced for cultures on the crop residue media, furfural media, and zero carbon medium, were reported. (Numbers in parentheses are the subset genes that are **exclusively expressed** and **exclusively silenced** in cultures on furfural media).
<sup>5</sup> The light-exposure experiment tested dark and four light-exposure timepoints: 15-, 60-, 120-, and 240-minutes reporting expression of 9728 genes. The number of genes expressed or silenced in the light or in the dark are reported (numbers in parentheses are the subset genes that are **exclusively expressed** or **exclusively silenced**).

A total of 68 *Neurospora* genes with HET domains (HET-domain genes) were identified and mapped to the *N. crassa* FGSC2489 strain [Figure 1 in reference 33]. Many of these HET-domain genes exhibited nonrandom distributions and were clustered near the end of the linkage groups, largely overlapping with clusters of LSGs (**Fig. 2**). A previous study demonstrated that another HET-domain gene, NCU03125 (*het-C*), plays a role in vegetative incompatibility [37]. 42 of the 69 HET-domain genes were clustered with at least one LSG within a range of five genes (`max-gap = 5`), and 23 and 14 HET-domain genes were clustered with at least one LSG within a range of three (`max-gap = 1`) or two genes (`max-gap = 0`) separately (**Fig. 2A–G, Tables 1** and **S4**). In fact, many of HET-domain genes were only syntenic within *Neurospora* and very closely related species in the Sordariales, and most HET-domain genes were surrounded by *Neurospora* LSGs and comparatively “young” genes (**Fig. 2H, Table S5**).

### Many LSGs and HET-domain genes are dynamically expressed in response to developmental and environmental changes

Genome-wide gene expression was measured in key stages of the *N. crassa* life cycle. LSGs and non-LSG genes generally exhibited substantial differences in the numbers of genes that were effectively expressed and silenced (**Figs. 1**, **S2, S3**; **Tables 2, S6**). During sexual development on Synthetic Crossing Medium (SCM) and asexual growth on Bird Medium (BM), similar trends were observed for proportions of LSGs and non-LSG genes that exhibited measurable expression, but an increased proportion of LSGs were expressed during the early hyphal branching on Maple Sap Medium (MSM; **Fig. S2**). At one or more time points during perithecial development, 238 genes exhibited at least 5-fold (*P* < 0.05) expression changes during perithecial development, indicating potential roles in the regulation of sexual reproduction. 35 genes exhibited no measurable expression in any sampled life history stages, suggesting either function only in unusual circumstances or mis-annotation as expressed genes (**Table S6**). During conidial germination and early asexual growth, respectively, 95 and 137 LSGs exhibited at least 5-fold (*P* < 0.05) expression changes for cultures on BM and MSM, and 213 and 181LSGs were expressed during none or only one of the four sampled stages for cultures on BM or MS separately. Expression of 342 LSGs was detected in at least two sampled stages during sexual reproduction and during asexual growth on asexual specific BM and MSM (**Fig. 3**).

**Figure 3.**
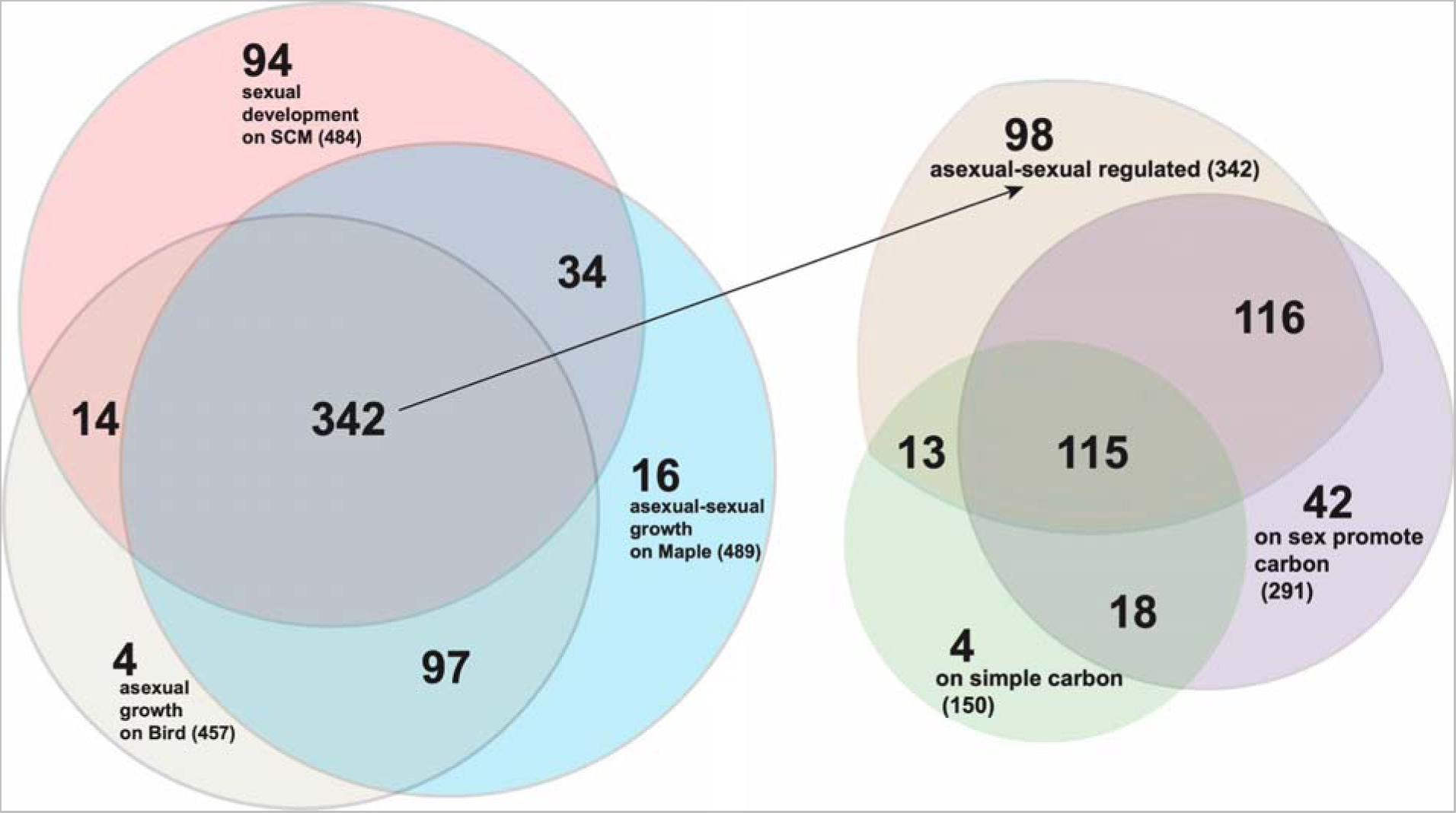
Differential expression of LSGs in *N. crassa* growth during three developmental processes [22,23] and on several media each supplying distinct carbon resources [39,40]. 342 LSGs were expressed at measurable levels in at least two stages in each of the three developmental processes (**Tables S6–S7**), including eight stages of sexual development on SCM (salmon pink, totaling 484 LSGs with measurable expression), four stages of asexual growth on BM (beige, totaling 457 LSGs with measurable expression) and four stages of asexual-sexual growth on MSM (light blue, totaling 489 LSGs with measurable expression). Among the 342 LSGs (arrow-linked), 231 were detectably expressed when cultured on either 2-furaldehyde furfural and/or 5-hydroxymethyl furfural (HMF), substrates that promote sexual development (purple, total 291 LSGs); and 128 were detectably expressed on sucrose and/or residues of at least one of five common crop straws (barley, corn, rice, soybean, and wheat; **Tables S6–S7**), substrates that support asexual growth and sporulation (light green, totaling 150 LSGs).

Genomic gene expression was also assayed for *N. crassa* growing on seven different carbon conditions including, absence of carbon, solo glucose carbon source, and a complex carbon source with five commonly available crop residues, including barley, corn, rice, soybean, and wheat straws, found in the field [39]. Expression of 464 LSGs was too low to be detected under any of these conditions; this large portion of *Neurospora* LSGs is not required for vegetative growth associated with carbon metabolism regulation (**Tables S7, S8**). Expression of 22 LSGs required that at least one type of carbon resource was present in the media, while expression of 56 other LSGs was only detected in the absence of carbon. Revisiting expression data collected from experiments investigating *N. crassa*’s tolerance to furfural [40] identified that 245, 239, 257, and 232 LSGs exhibited measurable levels of expression in the simple carbon cultures, DMSO (the carbon blank control), furfural, and HMF treatments (**Tables S7–S8**).

Within the 291 genes expressed in either furfural or HMF cultures or both, 61 were not expressed in the wild-type condition. Furfural is derived from lignocellulosic biomass and enriched in a post-fire environment. *N. crassa* sexual spore germination can be induced by furfural presence [41,42]. Furfural also inhibits conidia germination. Compared with cultures under wild-type conditions, 12 LSGs exhibited a 3-fold or higher expression in response to furfural, with NCU09604, 07323, and 01153 showing 6- to 22-fold up-regulation in furfural cultures.

Environmental factors, including light and temperature that regulate fungal growth and development, also dramatically affect expression of LSGs (Table S8 for more details). *N. crassa* genome is equipped with a set of light sensors responding to different light spectrums, duration and intensities [43–48], and *Neurospora* LSGs exhibit sensitive responses to light conditions.

When *N. crassa* cultures were exposed to light up to 4 h, genes were classified as short light responsive genes or long light responsive genes based on their expression profile changes [46]. Revisiting the previous data disclosed that out of 488 genes induced by light stimulus, 106 were LSGs (significantly enriched, *P* < 0.01), 59 of which were in the predicted clusters, including all genes in three 2-LSGs clusters, including cluster #25, 61, and 81 as well as three genes in a 4-gene cluster #127. Among 49 genes whose expression halted upon exposure to light, six were LSGs, including two genes NCU05052 and 05058 in a 3-gene cluster (**Fig. 4A, Table S9**).

**Figure 4.**
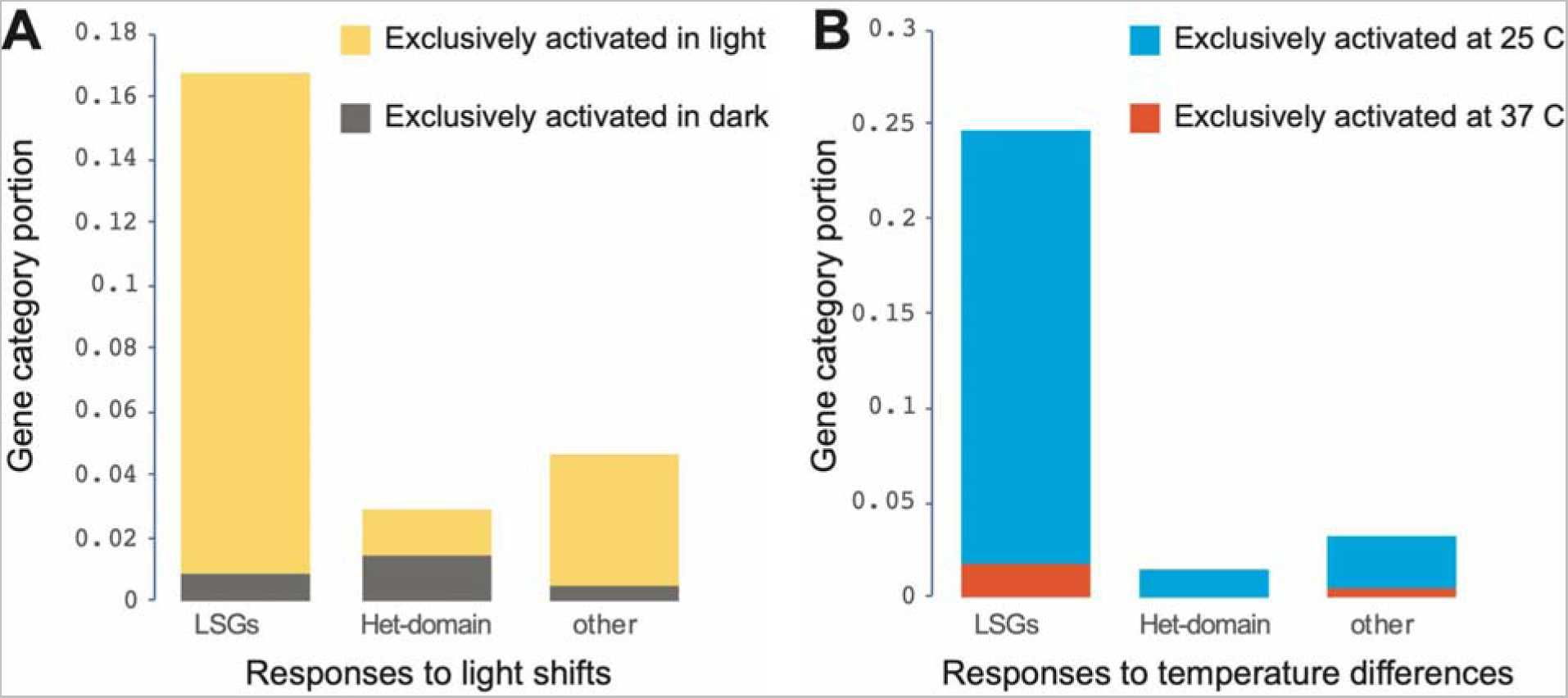
Genome-wide expression responses of LSGs, HET-domain, and other genes to shifting of culture environment. (A) Stacked proportions of 670 LSGs, 68 HET-domain, and other genes in the *N. crassa* genome, exhibited expression only in the dark (gray) or only in response to 15 to 240 minutes of light exposure (yellow). (B) Stacked proportions of 670 LSGs genes, 68 HET-domain genes, and other non-LSG genes in the *N. crassa* genome that are expressed across 4 stages of conidial germination and vegetative growth only at 25 C (blue) or 37 C (red) on Bird medium.

During conidial germination at a high temperature of 37 C on BM, expression of 270 LSGs was completely inhibited, a substantial enrichment (*P* < 0.01; a total of 941 out of 10592 genes exhibited no measurable expression). There were 148 LSGs exhibiting no measurable expression at 25 C and 37 C. There were 152 LSGs exhibiting detectable expression in cultures at 25 C but being turned off at 37 C, 100 of which were clustered LSGs, including 24 clusters with more than 2 genes that were turned off at 37 C. In contrast, 13 genes exhibited detectable expression in cultures at 37 C, but not at 25 C (**Fig. 4B, Table S9**). The LSG cluster #25 of NCU02144–02145 was the only LSG cluster that was silent in dark conditions or at 37 C.

Expression of HET-domain genes as a whole exhibited no clear patterns in response to environmental conditions or developmental stages **(Fig. 4A–B, Tables S8–9**). However, 11 HET-domain genes were not expressed in cultures on furfural or HMF, and 28 HET-domain genes were expressed neither in the dark nor during a shift from the dark to light for a duration of up to 2 h. Both of these conditions are known to promote sexual reproduction in *N. crassa*. More genes () were significantly up-regulated (5968 vs. 2935; *P* < 0.05) during the first branching of the germ tube on MSM, which supports both asexual and sexual development, than those on BM, which is specifically designed for promoting asexual reproduction and inhibiting sexual development. Accordingly, many more HET-domain genes were significantly up-regulated (46 vs. 15; *P* < 0.05) on MSM than on BM during that first branching stage.

### Some clustered LSGs and HET-domain genes exhibit coordinate expression

The majority of LSGs were clustered into physically linked groups—64% with *het* or HET-domain genes (Table S4). Forty-two HET-domain genes clustered with at least one LSG, including HET-domain LSGs NCU03378, 07596, and 10839. Other than these three HET-domain LSGs, all clustered HET-domain genes exhibited measurable expression in at least three out of the four sampled stages in conidia germination as well as six out of eight stages sampled during sexual development.

Coordinated expression among genes within the clusters during *N. crassa* asexual and sexual growth and development was not common. Among 26 cases where gene-expression dynamics were highly coordinated (at least one of the pairwise correlation coefficients were greater than 0.5 among LSGs in the cluster; **Table S10**) across several developmental stages, two are notable (**Fig. 5**). The first is cluster #69 of *het-14* and two LSGs (NCU07510 and 08191).

**Figure 5.**
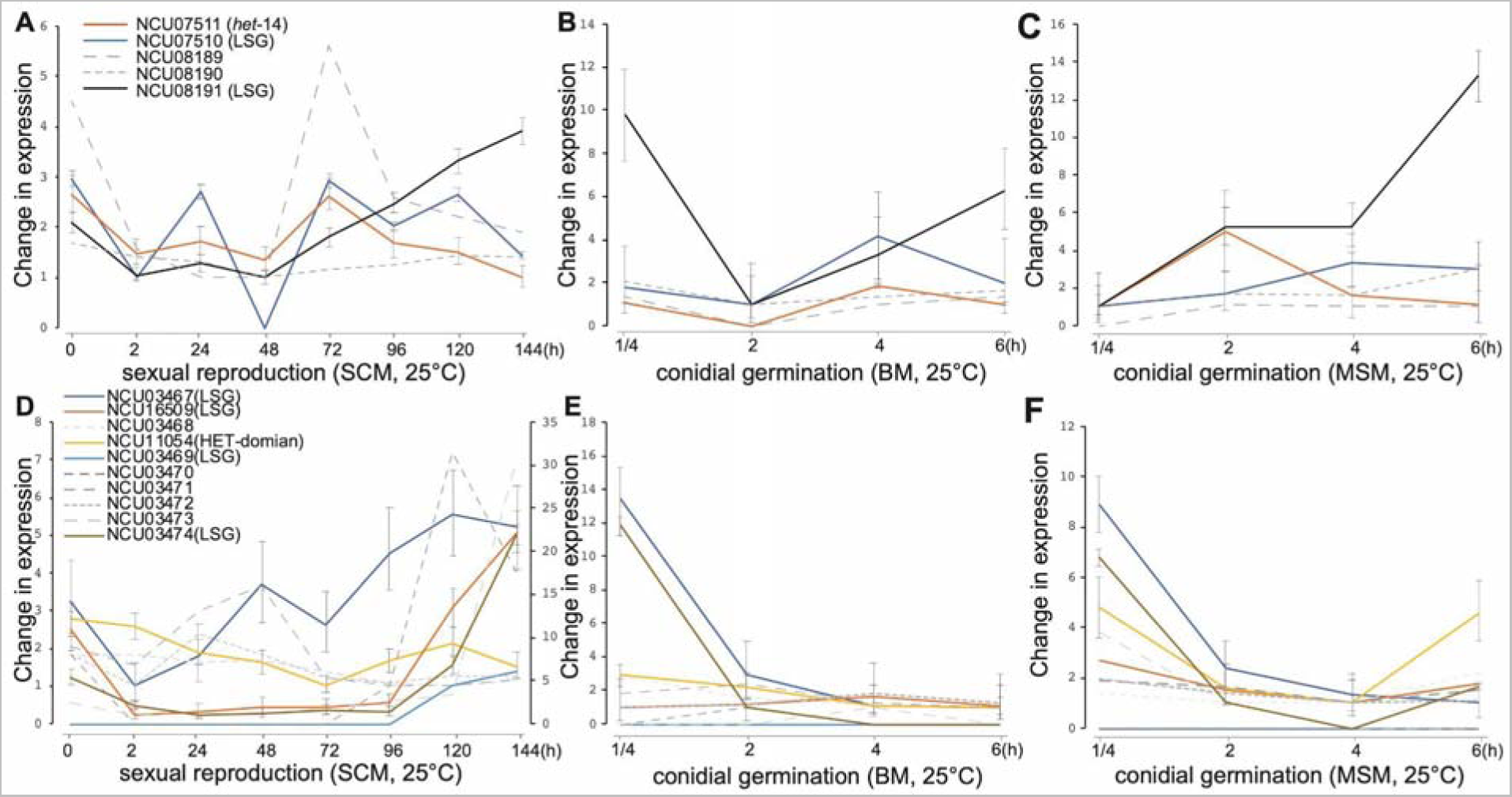
Expression profiles of two LSG-het gene clusters across asexual and sexual growth in *N. crassa*. expression profiles are plotted for LSG and HET-domain genes clustered with het-14 (color coded, with 95% credible intervals)—as well as the non-LSG genes NCU08189 and NCU08910 located within the cluster (grey dashed)—cultured through (A) eight stages of sexual development on synthetic crossing medium, (B) four stages of conidial germination and asexual growth on Bird medium, and (C) four stages of conidial germination and asexual growth on maple sap medium. Expression profiles for genes clustered with het-14 (color coded, with 95% credible intervals), including the five non-LSG genes NCU03468, 03470, 03471, 03472, and 03473 (grey dashed) within the cluster, across (D) eight stages of sexual development on synthetic crossing medium, (E) four stages of conidial germination and asexual growth on Bird medium, and (F) four stages of conidial germination and asexual growth on maple sap medium.

Genes in the cluster with *het-14* exhibited highly coordinated expression during sexual development, even with a large non-coding sequence of over 15000 bp separating *het-14* and NCU07510 in that cluster (**Fig. 5A–C**). Interestingly, the non-LSG genes located within the cluster exhibited expression dynamics that were similar to the clustered orphan and HET-domain genes during those stages. The second notable case where gene-expression dynamics were highly coordinated across several developmental stages is the cluster #117 of a HET-domain gene (NCU11054) and four LSGs (NCU03467, 03469, 03474, and 16509). Genes in the cluster #117 exhibited highly coordinated expression during asexual growth on MSM (**Fig. 5D–F**).

Expression was also observed to be coordinated in a few other clusters, such as among the LSGs clustered with the HET-domain genes NCU07335, 10142, and 16851 during conidial germination and asexual growth on BM and on MSM (**Fig. S4**): three clusters exhibited coordinated expression across sexual development (**Fig. S4A–C**), eight exhibited coordinated expression across conidial germination and asexual growth on Bird medium (**Fig. S4D–K**), and 15 exhibited coordinated expression across conidial germination and asexual growth on maple sap medium (**Fig. S4L–Z**).

Two of the three LSGs clusters, including the cluster #24 (**Fig. S4A**) and the cluster #131 (**Fig. S4C**), exhibited coordinated expression among the genes within each cluster during sexual development, and the coordinate expression was observed during the early stages of sexual development before meiosis (about 48–72 h after crossing). Of the eight clusters, expression during asexual growth on BM was coordinately down-regulated in seven of them (**Fig. S4D–K**). The exception was cluster #50 of HET-domain NCU07335 and LSG 07336; expression of these genes was up-regulated during sexual development (**Fig. S4I**). Coordinate expression during asexual growth on MSM exhibited an opposite pattern: 12 out of 15 clusters exhibited up- regulated expression patterns toward the extension of the first hyphal branch (**Fig. S4L–Z**), including two LSG-*het* clusters: the cluster #50 (**Fig. S4S**) and the cluster #51 of NCU16851 (HET-domain), 07316, 07317, and 07323 (**Fig. S4T**). In fact, coordinate expression was not only observed between LSGs and HET-domain genes that were clustered together, but also observed between LSGs in the clusters and neighboring non-LSGs (**Table S10**). Furthermore, genes in cluster #112 exhibited no measurable expression in least at three out of the four sampled stages across conidial germination and across asexual growth, and genes in cluster #172 exhibited no measurable expression in at least six out of eight sampled stages in sexual development in *N. crassa*.

### Cell communication transcription factors affect expression of LSG and HET-domain genes

The transcription factors *adv-1* and *pp-1* play multiple roles in asexual and sexual development, cell growth and fusion, and cell communication in *N. crassa* and closely related fungi [49–53]. Functions and regulatory networks involving *adv-1* and *pp-1* were systematically investigated [53], and in that study 155 genes were identified that were likely positively regulated by both transcription factors, including two HET-domain genes (NCU03494 and 09954) and one LSG (NCU17044). We reanalyzed the RNA sequencing data collected during conidial germination from knockout mutants of *adv-1* and *pp-1* and from wild type strain [53]. In general, knocking out the two transcription factors had a substantial impact on the activities of LSGs (**Figs. 6, S5, Tables 3, S11, S12**). Unlike HET-domain genes and other non-LSG genes, a significantly large portion of 173 LSGs exhibited no expression in the knockout mutants and the wild type strain, including 196 not expressed in both Δ*pp-1* and the wild type, and 195 not expressed in both Δ*adv-1* and wild type. At the same time, significant numbers of LSGs were turned on or off by the mutations to the two transcription factors. Namely, the same number of 44 LSGs that were expressed in wild type were inactivated in the mutant strains Δ*pp-1* or Δ*adv-1*, and 23 LSGs were inactivated in both the mutant strains Δ*pp-1* and Δ*adv-1.* Among the 23 LSGs, 20 were within the predicted LSG-het clusters, including NCU04700 and 04710 that clustered with HET-domain gene NCU04694 and three core genes NCU08822, 08829, and 08830 in a six-gene cluster. At the same time, expression of significant numbers of LSGs (53 and 54 separately) went from undetectable to detectable in the Δ*pp-1* or Δ*adv-1* knockout strains, with expression of 31 LSGs becoming detectable in both knockout mutants. However, only 14 out of the 31 newly detectably expressed genes were clustered LSGs. Therefore, knocking out these two transcription factors did not positively affect the LSG-*het* gene clusters.

**Figure 6.**
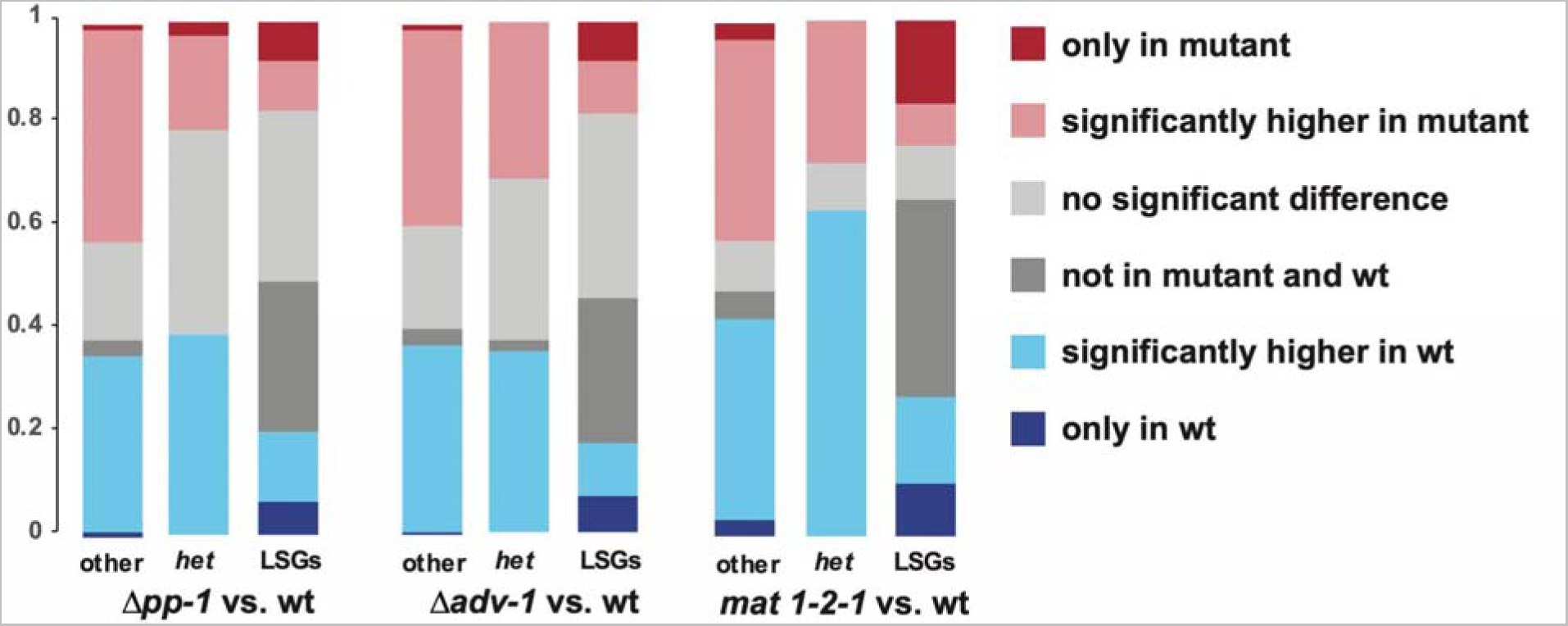
Impacts of transcription factor (TF) deletions on expression of HET-domain and LSGs within the *N. crassa* genome. Comparatively larger portions of LSGs are expressed exclusively in the mutants (dark red) or in the wildtype strains (dark blue). Expression profiles were classified in five categories (dark red: only measurable expression in mutant; light red: higher in mutant with *P* < 0.05; light grey: no significant difference with *P* ≥ 0.05; dark grey: not measurable in either mutant or wildtype; light blue: significantly higher in wildtype with P < 0.05; dark blue: only present in wildtype;), and comparative portions of each category in HET-domain genes, LSGs, and other genes in the genome were reported. Data from Fischer et al. (2018) were reanalyzed to assess the impacts of *pp-1* and *adv-1* knockout mutants.

**Table 3.**
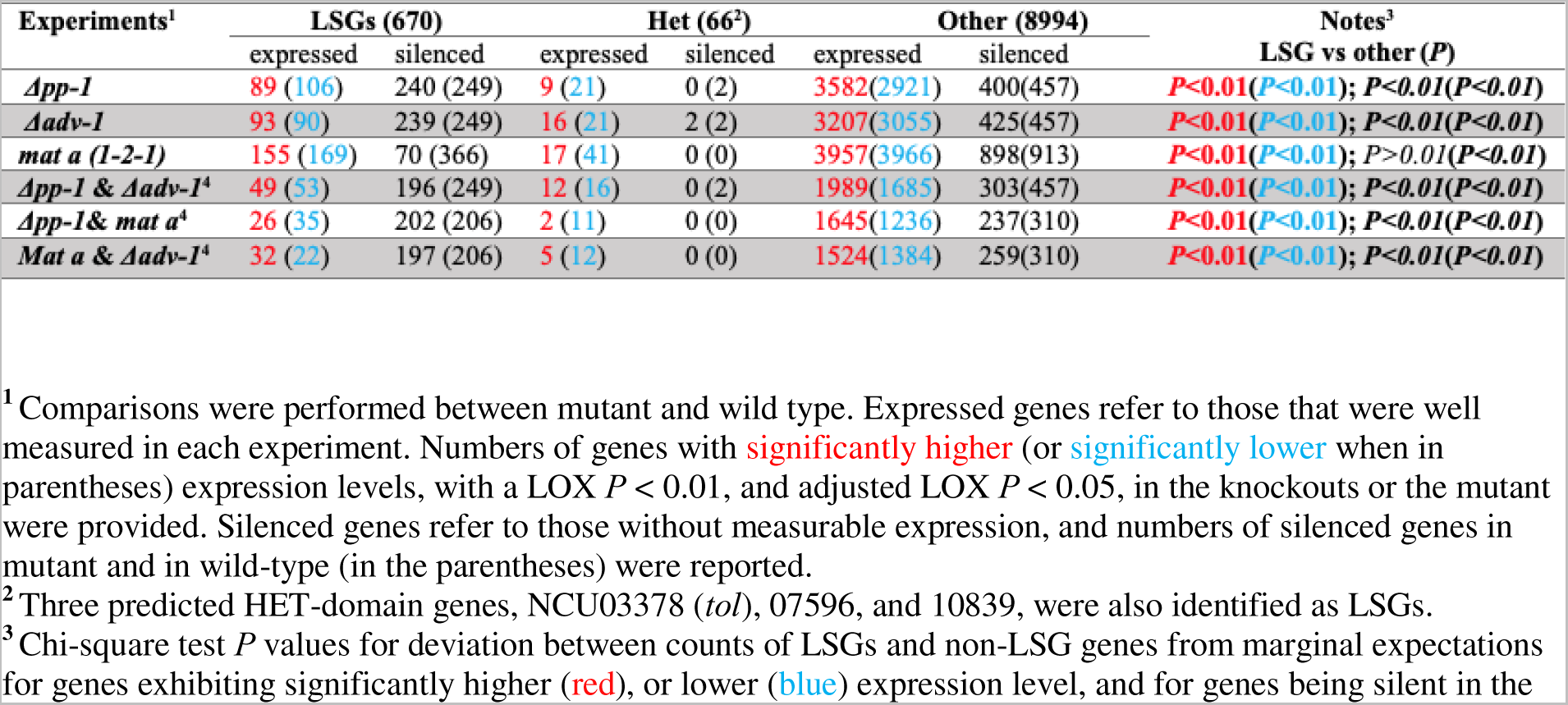
Transcriptomics analysis results on *N. crassa* responses to genetic manipulation.

Twenty-seven HET-domain genes are expressed at higher level in the wildtype strain than in both mutants (41 for Δ*pp-1*, and 35 for Δ*adv-1*), including four HET-domain genes (NCU03494, 06583, 09037, and 09954) exhibited more than five-fold higher expression in the wild type strain than in the Δ*pp-1* and/or Δ*adv-1* strains. However, NCU03494, 06583, and 09037 are HET-domain genes not clustered with any LSGs. For LSGs and HET-domain genes with well measured expression in mutants and wild type, many genes exhibited similar up- or down-regulation between the Δ*pp-1* and Δ*adv-1* compared with their expression in the wild type (**Fig. S5**).

*N. crassa* mating loci regulate crossing and sexual development, and opposite mating-type strains are ordinarily vegetatively heterokaryon-incompatible [54–57]. Transcriptome profiles were compared between six-day cultures of the wildtype strain and a mating locus *mat 1-2-1* mutant that has lost mating function (FGSC#4564, *mat a*[m1]s-3B cyh-1) on synthetic crossing medium (SCM). Of 9758 measured genes, a total of 836 genes exhibited undetectable expression only in either the mutant or wildtype (**Figs 6, S5**; **Tables S11, S12**), including 179 LSGs (109 expressed and 70 undetected in the *mat 1-2-1* mutant) and 657 non-LSG genes (336 expressed and 321 undetected in the *mat 1-2-1* mutant). For LSGs, lack of detectable expression occurring only in the wildtype or only in the mutant was significantly enriched (*P* < 0.00001, chi-squared test). Of the 109 LSGs that were exclusively expressed in the *mat 1-2-1* mutant, 71 were located within 55 predicted LSG-*het* clusters. Of the 70 LSGs that were only expressed in the wild type, 59 were located within 47 predicted LSG-*het* clusters. Only 17 clusters were common between the two groups, and larger clusters with more than three genes presented behaved differently between the *mat 1-2-1* mutant and wild type, with three genes of five-gene cluster #51 (NCU07306–07323) being inactivated in the mutant, while four genes of seven-gene cluster #124 (NCU05480–06949) and five genes of the nine-gene cluster #94 (MCU07144–07152) being activated in the mutant. Some two-gene clusters exhibited coordinated expression that was detectable only in the mutant or the wild type. There were 53 LSGs expressed significantly higher (*P* < 0.05) in the mating type mutant, and 111 expressed significantly higher in the wild type. However, mutations in the mating locus have no such impacts on basic activities for HET-domain genes, with 18 and 41 HET-domain genes being expressed significantly higher in the mating locus mutant, and 41 HET-domain genes being expressed significantly higher in the wild type (**Figs. 6, S5**).

Binding-site enrichment analysis using CiiiDER [58] identified no significant enrichment of binding sites for *mat 1-2-1*, *adv-1*, or *pp-1* in the upstream 5000 bp of LSGs that exhibited activity divergence in the mutants, specifying LSGs that whose expression was unchanged between the mutant and wildtype as the background. LSGs and HET-domain genes were not significantly enriched in genes that are potentially bound by *adv-1* or *pp-1*, based on data from DAP-seq [SRP133627 from 53]. However, knocking out a key transcription factor encoding gene, *ada-6* [59]—whose product regulates asexual and sexual growth—inhibited expression of 30 LSGs and promoted expression of 25 LSGs during conidiation and protoperithecial production. Therefore, a substantial number of LSGs are at least peripherally involved in those regulatory networks, abundant enough that they may play fine regulatory roles, and sparsely distributed enough to impact diverse regulatory pathways.

### Selection profiles of LSGs

an original study reported 135,000 SNPs in transcriptomic sequence from 48 individuals [60], yet a total of 1,086,579 SNPs is reported from just 26 *N. crassa* isolates whose transcriptomic sequence data are available in FungiDB. The discrepancy likely arises as a consequence of the exclusion of singletons from the original data [60], which were explicitly left unreported in the original paper, but were frequently observed (C. Ellison personal communication, July 13, 2020). 97.8% (655 out of 670) LSGs feature at least one non-synonymous SNP, which is moderately but statistically significantly higher (Fisher exact test, *P* < 0.01) than the 91.6% (8271 out of 9027) non-LSG genes, as revealed by the inclusive SNP data in FungiDB (**Fig. S6**). Unlike LSGs that generally encode less than 250 amino acids, most HET-domain genes encode more than 300. Non-synonymous SNPs were observed at similar frequencies between LSGs and HET-domain genes with similar gene length (**Fig. 7)**. Excluding singletons from the FungiDB data markedly reduced the number of SNPs—only eight *Neurospora* LSGs then featured SNPs (1.3%) compared to 4192 non-LSG genes (45.9%).

**Figure 7.**
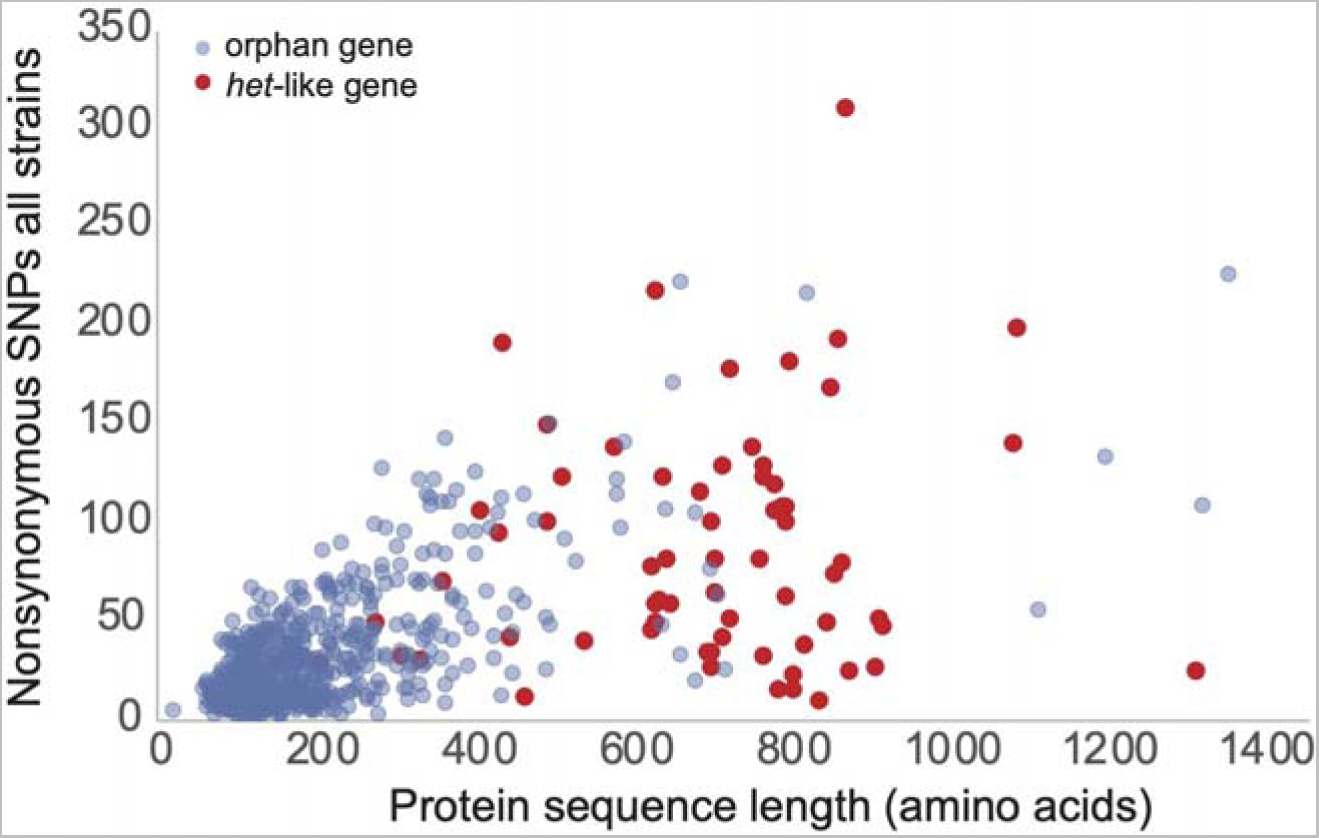
Nonsynonymous SNP distribution comparison between LSGs (blue circles) and HET-domain genes (red circles) from *N. crassa* population genetic sequencing [60]. Nonsynonymous SNPs are totaled across all 26 *N. crassa* strains deposited at FungiDB.

Therefore, SNP singletons are abundant within *Neurospora* LSGs. A high density of SNP singletons in short LSGs corresponds to a high allelic polymorphism of LSGs, present in a comparative low proportion in the population, implying high linkage disequilibrium in LSGs.

Of 603 *Neurospora* LSGs with SNPs data, including those with singleton SNPs, only NCU07210 exhibited statistically significant strong gene-regional positive selection during the divergence of *N. crassa* from its sister species *N. tetrasperma* (**Fig. S7A**). No abnormal phenotype resulting from the knockout of this gene (*mat-a* strain FGSC14291) was observed in our assays. For 695 LSGs shared within *Neurospora*-*Sordaria*, 16 genes exhibited statistically significant evidence of strong gene-regional positive selection, and five exhibited statistically significant evidence of moderately strong gene-regional positive selection (**Table S13**). Five of 16 strongly positively selected genes were identified in six biological processes related to the response to stress or stimulus (**Table S14**), including the hypothetical protein coding gene NCU02932 with the ortholog being annotated as a regulator of ATP-sensitive K^+^ channels in *N. discreta* and likely play roles in stress responses (**Fig. S7B**). Further exploration suggested that positively selected *Neurospora-Sordaria* LSGs NCU05395, 07618, 09562, and 09693 are likely involved in the regulation of sexual development [61]. Furthermore, regulation of the moderately positively selected genes NCU01306, 00496, and 00748 is likely associated with conidiation in *N. crassa* [59,62,63].

### KO phenotypes of LSGs

In an examination of phenotypes of crosses of 367 available KO strains of the *Neurospora* LSGs to the KO of the opposite mating type (or the WT of the opposite mating type if the KO of the opposite mating type was not available), two *Neurospora* LSGs, NCU00176 and 00529, and one *Neurospora*-*Sordaria* LSG, NCU00201, showed a distinct knockout phenotype in sexual development (**Fig. 8**). All these knockout mutants exhibited arrested development with the protoperithecia failing to develop into perithecia. Interestingly, both NCU00176 and NCU00201 were expressed at significantly higher levels in protoperithecia than in mycelia after crossing. Expression of NCU00529 was observed only in the late stage of conidial germination on maple medium. NCU00529, 00530, and 00531 are homologs, and NCU00529 and 00530 exhibited no expression in sexual development and conidial germination.

**Figure 8.**
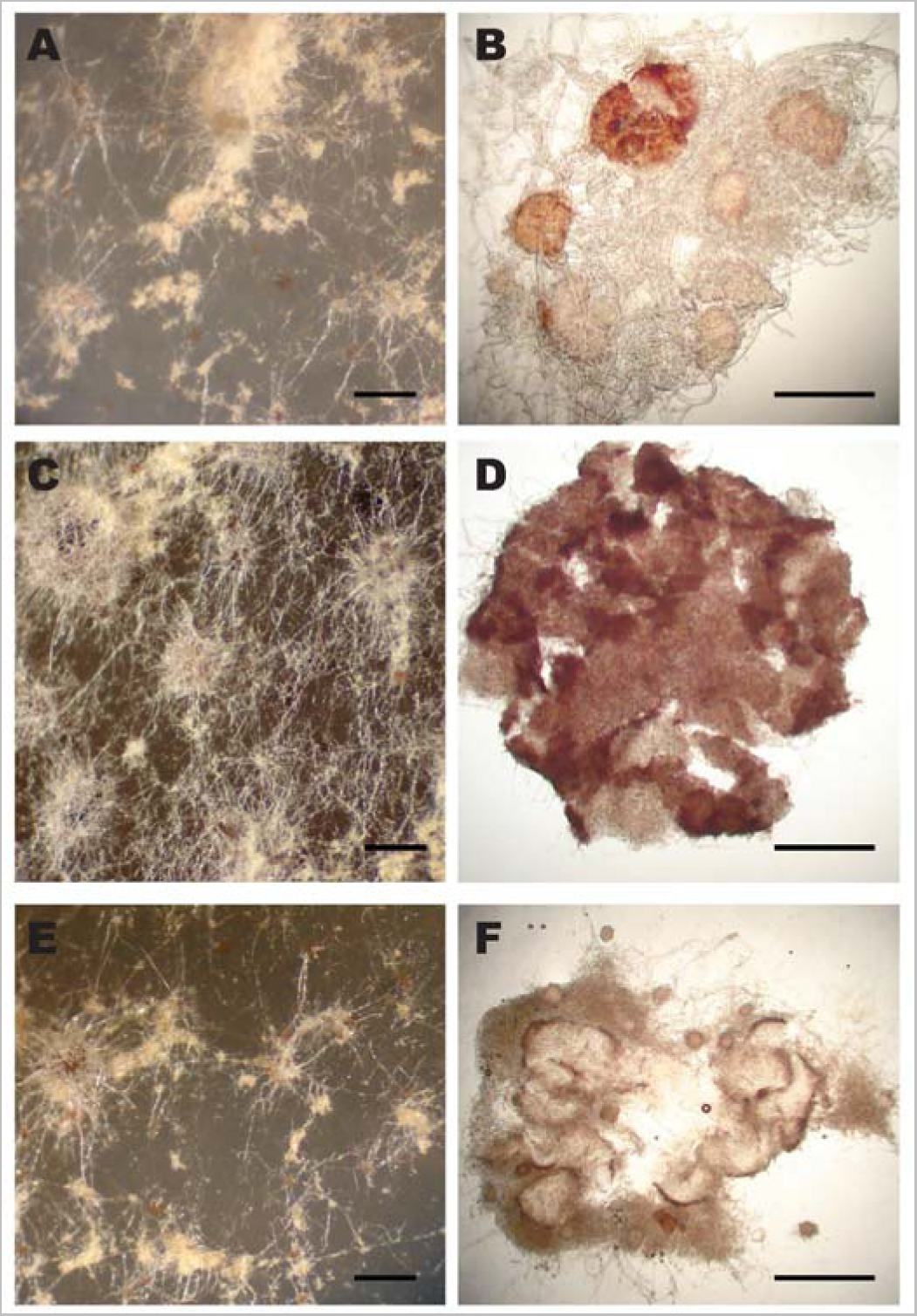
Knockout mutants of *Neurospora crassa* genes NCU00176, NCU00201, and NCU00529 produced arrested protoperithecia that failed to develop into perithecia and to produce sexual spores on SCM medium. Knockout cross ΔNCU00176 (FGSC12195 × 12196) exhibited (A) small protoperithecia (scale bar: 1 mm) and (B) squashed protoperithecia exhibiting an abortive ascogenous center (scale bar: 10 µm). Knockout cross ΔNCU00201 (FGSC18867 × 18868) exhibited (C) normal-sized protoperithecia (scale bar: 1 mm) with (D) abortive ascogenous centers (scale bar: 10 µm). Knockout cross ΔNCU00529 (FGSC13076 × 13077) exhibited (E) normal-sized protoperithecia with (F) abortive ascogenous centers.

No abnormal phenotypes were identified for NCU00531 knockout mutants (FGSC13078 and 13079), and no knockout mutants for NCU00530 are available. Knockouts of NCU00375, 00384, 00485, and 00491 (*Neurospora* LSGs) and NCU01623, 05395, 07618 (*Neurospora-Sordaria* LSGs) have been reported to exhibit minor phenotypic anomalies during asexual growth, especially at 37 C (Fungidb.org/fungidb). We examined knockout mutants of these genes and confirmed increased pigment production in NCU00485 and dense and slow growth in NCU00491 and 016223 at 37 C.

## Discussion

In this study, we systematically investigated LSGs in *Neurospora* genomes to determine their location and organization on chromosomes and possible roles in biology and ecology. Using genomic phylostratigraphy and reciprocal BLAST searching, we identified 670 genes that are *Neurospora* LSGs, only shared within *Neurospora* species. More than 63% of the 670 *Neurospora* LSGs form clusters of 2–21 LSGs, interspersed with HET-domain putative allorecognition genes. Many of the larger clusters aggregate near the telomeres of the seven chromosomes. A majority of *Neurospora* LSGs are actively regulated in response to carbon sources, light, and temperature changes that promote sexual or asexual reproduction. Knockout mutant strains were phenotyped for 367 *Neurospora* LSGs, and arrested protoperithecia were observed for mutants of three *Neurospora* LSGs. Our data indicate that regulation of asexual and sexual reproduction surprisingly empower evolution of *de novo* elements more than the regulation of core metabolism and cellular biology in *N. crassa*. More specifically, this regulation is associated with reproduction modes affected by carbon resources—a critical environmental factor for this post-fire fungus. However, many LSGs were not identified as essential for *N. crassa* development and biology. Therefore, it is possible that LSGs play key roles in fine regulation in response to environmental factors inducing reproduction processes that have been understudied.

### Neurospora LSGs and HET-domain genes are tightly clustered

*Neurospora* LSGs and HET-domain genes exhibit several organizational features on chromosomes. These organizational features include gene clustering, large non-coding spaces enclosed by flanking condensed coding regions, high frequencies of gene duplications, and proximity to telomeres. If clustered genes were derived from relocation and rearrangement, especially for functionally associated genes, it would probably be easy for these genes to functionally integrate back altogether into the existing system. Indeed, coordinated expression across clustered LSGs and HET-domain genes was sometimes detected—especially during *N. crassa* development in response to various environmental factors. However, LSG-*het* gene clusters and their neighbor genes were syntenic mainly within very closely related taxa, and many LSGs exhibited no expression under common laboratory settings—even across a range of nutrient conditions and developmental stages.

Interestingly, heterochromatic interactions of intra- and inter-telomeric contacts were reported as common in *N. crassa* [64], implying potential *cis-* and *trans-*regulation at chromosomal level of expression integration in these LSGs enriched regions. It is possible that integrating LSG-*het* gene clusters into existing regulatory systems temporally and spatially modifies the original functions that are expected only under common growth conditions. Therefore, specific experiments conducted in extreme growth conditions, with rare nutrient types, and at understudied stages of the life cycles should be conducted.

A previous study characterizing chromosome ends in *N. crassa* demonstrated that highly AT-rich sequences in the telomeres are likely products of RIP and that subtelomeric elements common in other fungi are absent in *N. crassa*. Telomere repeats are required for H3K27 methylation, which would repress the transcription activities and functionally silent genes in these regions [65]. More importantly, the telomeric regions have potential significance in niche adaptation and probably harbor hotspots for novel sequences due to abrupt sequence divergence involving repeats [66]. Many genes of an annotated *het*-domain also locate near the ends of some *N. crassa* chromosomes [33], and 42 of the 69 HET-domain proteins, which account for heterokaryon incompatibility, are actually clustered with at least one *Neurospora* LSG. In fact, neighboring regions of HET-domain genes are abundant with “young” genes that lack homologs in lineages beyond *Neurospora* and closely related species in the Sordariales, and most neighbor genes are syntenic within Sordariomycetes. Most LSGs were not active in sampled stages in the *N. crassa* life cycle. However, a few LSG-*het* gene clusters exhibited coordinate expression during early sexual development as well as during active hyphal tip growth on the maple sap medium that support both asexual and sexual developments. Therefore, investigation of gene activities in the telomeric and sub-telomeric regions during mycelium development and allorecognition between same and different vegetative compatibility groups will shed light on possible concerted associations between these two functional groups in heterokaryon incompatibility, pre-mating process and even during early sexual development in *Neurospora*.

Actually, three LSGs, including NCU03378, 07596, and 10839, were annotated as a *het*- domain, and NCU03378 was annotated as *tol* [tolerant, 55] and shared several conserved sequence regions with *het*-6 that is involved in incompatibility in the *N. crassa* population [67]. Functional associations among some HET-domain genes and LSGs within the LSG-*het* clusters were evidenced with similar gene expression regulation pattern during asexual and sexual development in *N. crassa*. However, compared with the HET-domain genes that were all expressed, many LSGs were not expressed in the sampled stages during the *N. crassa* life cycle. Further investigation on epigenetic modification of gene expression near telomeric neighborhoods could be critical to capture the right moments of associated functions between LSGs and HET-domai*n* genes during allorecognition and to understand the evolution of LSGs in *N. crassa*.

### Neurospora LSGs play roles in response to key regulatory environmental factors

LSGs are generally not functionally annotated, mainly due to the lack of homologous references in well studied genomes. To determine the functional roles of *Neurospora* LSGs, we revisited recent high quality transcriptomics studies on *N. crassa*, covering almost all morphological stages in the *N. crassa* life cycle, cultures under different conditions and carbon or nitrogen resources, light exposure, temperature change, as well as knockout mutants of key regulatory genes [22– 24,39,40,46,59,68–72]. We observed a significant enrichment of clustering of LSGs and HET-domain genes and, within some of those clusters, highly coordinated regulation in response to carbon resources, light, and temperature conditions. This study provides evidence that some LSGs and clustered HET-domain genes are associated with metabolic adaptation to environmental factors that are critical indicators of successful fungal asexual and sexual growth and reproduction.

Three hundred forty-two LSGs that were actively expressed during sexual development were also actively regulated in both asexual growth on two different media, including Bird medium, which supports only asexual reproduction, and maple sap medium, which supports both asexual and sexual growth. Nearly two thirds of the 342 genes were actively regulated in samples collected from the media supplied with carbon resources that promote sexual growth of *N. crassa*, supporting their possible roles in sexual development and the asexual-sexual switch.

In favor of their roles in sexual development in *N. crassa*, more than one third of the LSGs were actively regulated on the presence of specific carbohydrates, including HMF and furfural, compounds that powerfully stimulate initiation of sexual reproduction. *N. crassa* has been shown to respond differentially to the two furans and to possess a high tolerance to furfural, which is present in its natural habitat [40]. It is conceivable that some of the LSGs that are uniquely expressed upon exposure to HMF or furfural could be further engineered to provide increased tolerance to atypical carbon resources for *N. crassa*, a trait of significant interest in the pursuit of robust biofuel production.

*Neurospora* LSGs were observed to be active only at the hyphal tips without measurable expression from the colony samples, and thus these genes were considered to be responsible for environmental sensing and interaction with microbes [73]. Overall, LSGs are probably associated with reproductive development and growth in response to environmental conditions, especially carbon resources, light and temperature. Furthermore, we observed that LSGs in predicted clusters coordinately respond to these environmental factors. How these genes may possibly contribute together as fine quantitative switches for reproduction requires further investigation with multiple gene manipulations.

### A significant proportion of LSGs are regulated by key developmental transcription factors

Transcription factors, including *pp-1* and *adv-1* that play key roles in cellular communication during asexual growth and mating loci that regulate sexual crossing, have been previously reported [53,54,74]. From the previous transcriptomics data from knockout mutants of *pp-1* and *adv-1* and newly generated transcriptomics data from a mutant of *mat 1-2-1*, we observed that the expression of a significant portion of LSGs was affected by mutations in these transcription factors. Interestingly, over 95% of LSGs that were turned off in both *pp-1* and *adv-1* mutants belonged to predicted clusters, but less than 50% LSGs that were turned on in both mutants belonged to predicted clusters. The expression of six genes in the NCU07144–07152 LSGs cluster was actively regulated during sexual development, in knockout mutants of *adv-1* and *pp-1* that regulate cell communication with knockout phenotypes in sexual development [49,52], as well as in samples in the absence of carbon. Expression of some LSGs was also affected in knockout mutants of other regulatory genes, such as *ada-6* and *gul-1* that are critical for sexual and asexual development in *N. crassa*. The most dramatic impacts to the expression of LSGs were from the mutation in the mating locus, suggesting likely functional associations of the mating process and sexual development initiation for LSGs. However, no binding sites of transcription factors were observed enriched in the promote and up-stream sequences of LSGs being turned on or off by those factors, and coordinate expression regulation was only detected in a few LSG-*het* gene clusters. Instead of possible cis or trans regulations, an alternative explanation for the associations between the LSGs and transcription factors would be that the LSGs were coordinated with the sampled developmental stages when the transcription factors were actively engaged. For example, the LSGs exhibited different expression activities during hyphal branching on natural medium MSM, when cell-to-cell communication is regulated by *adv-1*. Therefore, success in further investigations will be aided by focusing on the epigenetics of the LSG-*het* gene clusters during specific periods of cell-to-cell communication.

### Knockout phenotypes suggested that few LSGs play essential roles in Neurospora development

Our phenotyping of available knockout mutants of 367 LSGs at the Fungal Genetics Stock Center [75] yielded three genes with a phenotype in sexual development. Within these three genes, NCU00529 and its homologs NCU00530 and 00531 form a cluster in the subtelomere of chromosome I. The other two genes with mutant phenotypes in sexual development, NCU00176 and 00201, exist on chromosome III. Their expression decreased after protoperithecia. NCU00176 was reported as one of the eight *Neurospora* LSGs that were among the 300 down-regulated hypothetical proteins in the *gul-1* knockout mutant [76], and GUL-1 plays multiple roles in *N. crassa* hyphal morphology development [77]. We also confirmed increased pigment production in NCU00485 and dense and slow growth in NCU00491 and 016223 at 37 C.

A recent study reported 40 biologically relevant clusters (BRCs) for 1168 *N. crassa* genes being phenotyped for 10 growth and developmental processes [78]. Eleven *Neurospora* and 31 *Neurospora-Sordaria* LSGs were included in this analysis. Interestingly, 4 out of the 11 *Neurospora* (*P* < 0.05) and 5 out of the 31 *Neurospora-Sordaria* LSGs were concentrated in one of the 40 BRCs (Cluster 4 of 81 genes) with minimum yeast orthologs and generally no significant phenotypes. An explanation for the lack of apparent phenotypes may be that these genes 1) were not investigated under the conditions for the phenotypes; 2) were functionally non-essential and/or not fully integrated into the regulatory networks, or 3) were functionally redundant within clusters or with non-LSG paralogs.

### LSGs are clustered with allorecognition loci and respond to G × E factors regulating the switch from asexual to sexual reproduction in Neurospora

The co-location of LSGs and HET-domain genes may suggest that they are functionally and evolutionarily linked. Our observations show that the expression of LSGs is significantly altered during the transition from asexual to sexual reproduction in *N. crassa*, in response to environmental factors such as carbon resources, light, and temperature, as well as genetic regulators such as cell-to-cell communication transcription factors and mating types. A substantial proportion of the LSGs are involved in inducing and promoting sexual development in response to G × E factors. Coincidentally, during the asexual-sexual transition, allorecognition governed by HET-domain genes, particularly mating loci, shifts from being active to being repressed, allowing for crossing between different Vegetative Compatibility groups (VCGs). Direct functional interactions between the two gene groups would be difficult to investigate using standard transcriptomics approaches, as allorecognition, which negatively regulates fusion between different VCGs, and sexual reproduction occur in different developmental phases and under different environmental conditions. Despite this, we observed limited coordinated responses to environmental and genetic regulatory factors. Further investigation at the species and population levels, with a focus on VCGs, is required to determine the evolutionary histories of LSGs and HET-domain genes, which are linked by their co-location in recent lineages. Therefore, SNP data from various populations, especially from different VCGs, would be of great interest in understanding the evolutionary associations between the two gene groups. In fact, balancing selection has been demonstrated to act on HET-domain genes, and abundant rare singletons have been observed for LSGs, with a similar occurrence of SNP polymorphisms between HET-domain genes and LSGs within the same molecular size, based on data from limited *N. crassa* populations

### A working model provides a foundation for further investigation of G × E interactions of LSGs and HET-domain genes in Neurospora reproduction

The discovery of the co-location of the LSGs and HET-domain genes at the fast-evolving hotspot in the *N. crassa* genome and of the LSGs participation in G × E interactions that regulate the asexual-sexual switch inspired us to develop a model to guide future investigation about possible interactions between LSGs and HET-domain genes that are challenging in a standard experimental setting for *N. crassa* molecular genetics (**Fig. 9**). The proposed working model is substantiated by the responses of LSGs and clustered HET-domain genes to environmental factors, developmental stages, and key regulatory transcription factors. This model suggests that the G × E regulations from asexual growth and reproduction shift to sexual reproduction in *N. crassa* connect to activities of both the LSGs and HET-domain genes.

**Figure 9.**
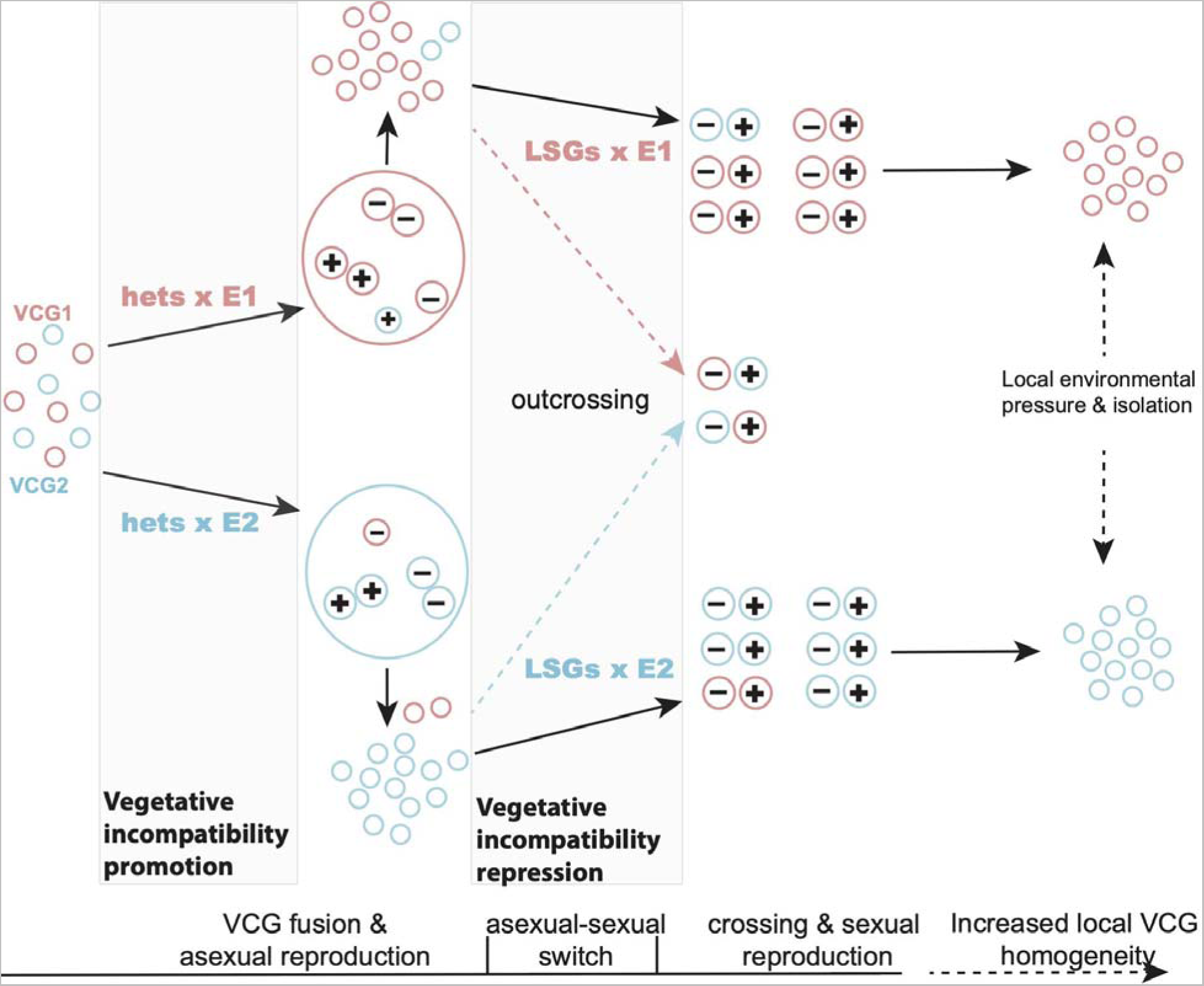
Hypothesis regarding the roles of LSGs associated with allorecognition and environmental regulation of reproduction modes. Pink and blue coded for vegetative compatibility groups (VCGs) that are governed by allorecognition (HET-domain genes), and regulated by the LSGs and environment for being adaptive to different growth environments E1 and E2 and are incompatible to each other. VCG colonies expand via aggregation and anastomosis under preferred vegetative growth conditions until the environment turns on the asexual-sexual switch and shifts into sexual reproduction. Plus (+) and minus (–) represent two opposite mating types. Crossing of closely-related strains would further promote local segregation of the VCGs, while outbreeding would contribute to polymorphism in the populations. LSGs may contribute to the consolidation of vegetative incompatibility when environmental conditions are preferred for asexual growth, as a few HET-domain genes are LSGs. LSG expression in environmental and genetic manipulations is consistent with roles in the induction of sexual development, as well as roles in the repression of vegetative incompatibility when environmental conditions are preferred for sexual reproduction. However, there is a lack of investigation of the complex but interlocking interactions among environment, LSGs, and VCGs, and their epistatic interactions during fungal asexual and sexual growth. Therefore, future investigations will reveal whether mutations in LSGs and allorecognition could contribute to parapatric or even sympatric speciation due to ecological selection, or characterize the accumulation of mutations in LSGs and HET-domain genes during allopatric and peripatric speciation due to physical barriers.

During asexual growth and reproduction, HET-domain genes restrict cell fusion to be successful only between mycelia of the same VCG groups. Studies of fungal pathogens on plants and animals suggest that fast-evolving LSGs, often as clusters, contribute to extensive chromosomal structure polymorphisms that may drive the aggressiveness during host colonization and the evolution of virulence [79–82]. When environmental conditions are favorable for sexual development, LSGs may react and play roles in the initiation of sexual development and the repression of HET-domain genes. Consistent with our hypothesis, introgression of *N. crassa het-6* gene (previously identified as *tol*) into *N. tetrasperma* disrupts the tolerance of both mating types in asexual tissues, thwarting pseudo-homothallism and causing death of *N. tetrasperma* [83]. Three HET-domain genes are actually LSGs in *N. crassa*. Nevertheless, the model proposes LSGs-environment associated mechanisms subsequent to vegetative incompatibility during asexual colonization that enable crossing between compatible mating types from different VCG backgrounds (**Fig. 9**).

This crossing between compatible mating types from different VCG backgrounds would induce local homogeneity of VCG, permitting increased colony merging, nutritional exchange, propagation of signals of disease and predators, and would increase the opportunity for mitotic recombination via parasexuality. There are no laboratory studies evaluating fitness of VCGs on controlled genetic backgrounds in different environments providing substantial evidence of differentially adapted VCGs. However, field studies in *Aspergillus* demonstrated that specific VCGs can have competitive advantages in specific agro-ecological zones [84]. Furthermore, *A. falvus* VCG groups isolated from cotton in the southern USA are genetically isolated, despite presumed sexual reproduction between the two opposite mating types, each of which was only detected in one VCG group [85]. Moreover, locally predominant VCGs have been considered crucial to the formation of isolated, asexual, inbreeding populations in *Fusarium oxysporum*, a species in which sexual stage and meiotic recombination have been lost, based on an evolutionary model proposed by Puhalla that was later supported by multiple studies [86,87]. Therefore, our model focusing on LSGs calls for investigation of how vegetative incompatibility that promotes the local aggregation and anastomosis of identical VCGs would be associated with specific environmental preferences in nutrients, temperature, light, and osmosis, as well as parasite evasion, or other competitive advantages. Our model of LSG G × E interaction also provides a route for experimental investigation of why there are predominant VCGs and rare VCGs in distinct geographic regions or environmental settings.

It would be also interesting to test if such purifying selection would lead to genomic homogeneity in allorecognition loci and be further enhanced supposing nearby crossings and sexual reproduction. Similar with lineage-specific genes in other model systems [e.g. 88,89], highly frequent SNPs, including SNP singletons, were observed in *N. crassa* LSGs, implying either weak selection, or recent selection, or both. However, SNP and SNP singletons, often considered to be at linkage disequilibrium, can be affected by various factors, and SNP singletons have been assigned functional importance in the human genome [90]. However, filtered SNP data from limited *N. crassa* population in FungiDB could not provide a full picture regarding population structures of LSGs. Therefore, our model also suggests that long term and large scale pangenomics would be critical to understanding whether outbreeding between VCGs plays a substantial role in promoting allelic diversity in *Neurospora* (**Fig. 9**).

## Materials and Methods

### Mutant and culture conditions

Protoperithecia were sampled for *mat 1-2-1* mutant (FGSC#4564, *mat a*[m1]s-3B cyh-1). The experiments were performed with macroconidia, which were harvested from 5-day cultures on Bird medium (BM). 1 × 10^5^ spores were placed onto the surface of a cellophane-covered synthetic crossing medium (SCM) in Petri dishes (60 mm, Falcon, Ref. 351007). Dark-colored protoperithecia were abundantly ripen in 6 days after inoculation. Tissue samples were flash frozen in liquid nitrogen and stored at -80LJ.

Biological replicates included all tissues collected from multiple plates in one collection process. Three biological replicates were prepared for each sampled point.

### RNA isolation and transcriptome profiling, data acquisition and analysis

Total RNA was extracted from homogenized tissue with TRI REAGENT (Molecular Research Center) as in Clark et al. (2008)[91], and sample preparation and sequencing followed our previous works [23,25,92]. Briefly, mRNA was purified using Dynabeads oligo(dT) magnetic separation (Invitrogen). RNAseq Library Prep: mRNA was purified from approximately 200 ng of total RNA with oligo-dT beads and sheared by incubation at 94 C in the presence of Mg (Roche Kapa mRNA Hyper Prep Catalog # KR1352).

Following first-strand cDNAtA-tailing was performed with dUTP to generate strand-specific sequencing libraries. Indexed libraries were quantified by qRT-PCR using a commercially available kit (Roche KAPA Biosystems Cat# KK4854). The quality of cDNA samples was verified with a bioanalyzer (Agilent Technologies 2100).

The cDNA samples were sequenced at the Yale Center for Genomics Analysis (YCGA). The libraries underwent 76-bp single-end sequencing using an Illumina NovaSeq 6000 (S4 flow cell) according to Illumina protocols. Adapter sequences, empty reads, and low-quality sequences were removed. Trimmed reads were aligned to the *N. crassa* OR74A v12 genome from the Broad Institute [93] using HISAT2 v2.1, indicating that reads correspond to the reverse complement of the transcripts and reporting alignments tailored for transcript assemblers.

Alignments with a quality score below 20 were excluded from further analysis. Reads were counted for each gene with StringTie v1.3.3 and the Python script prepDE.py provided in the package. StringTie was limited to report reads that matched the reference annotation. Sequence data and experiment details were made available (GSE199259) at the GEO database (https://www.ncbi.nlm.nih.gov/geo/).

Statistical analysis of the sequenced cDNA tallies for each sample was performed with LOX v1.6 [94], ignoring raw reads that mapped ambiguously or to multiple loci.

### Identification and verification of Neurospora crassa lineage specific genes (LSGs)

We applied a two-step strategy to identify and verify *Neurospora* LSGs, including (1) a phylostratigraphic approach to reveal putative LSGs and (2) confirmation using BLAST against all genomes available at NCBI and fungal genomes at FungiDB. In the first step, we employed previously published genomic phylostratigraphy for the *N. crassa* genome that reported over 2000 *N. crassa* orphan genes, discovered by genomic comparisons versus *Chaetomium globosum*, *Ascrospora strigata*, *Saccharomyces cerevisiae*, *Phanerochaete chrysosporium*, *Drosophila melanogaster*, and *Arabidopsis thaliana* [29]. *N. crassa* orphan-gene status was determined via the Smith-Waterman pairwise similarity of protein-coding sequences [95,96]. Classifications of genes in *N. crassa* were constructed as mutually exclusive groups ranked in by their phylostratigraphy, including Euk/Prok-core, Dikarya-core, Ascomycota-core, Pezizomycotina-specific, *N. crassa*-orphans, and others [11,97]. Putative *Neurospora* LSGs were also compared with the newest annotation of the *N. crassa* genome. Consequently, the number of predicted *N. crassa* LSGs based on the representative genomes was narrowed down to 1872 genes.

Many fungal genomes were recently published due to the efforts of the 1000 fungal genome project launched by the DOE Joint Genome Institute [32]. These fungal genomes— especially those well-sampled among closely related species—make it possible to be confident that LSGs are likely the product of de novo gene evolution rather than birth-and-death processes [98]. To verify that the 1872 genes are not present in species that were not analyzed within the phylostratigraphic and previous BLAST analyses, we employed BLASTp and tBLASTx to search in the entire NCBI GenBank database specifying an *E*-value cutoff of 0.05. To utilize newer genome annotations that were available on GenBank, BLASTp and tBLASTx were used again in FungiDB with an *E*-value cutoff of 10. These searches included genomes closely related to the Neurospora genomes, including *Podospora*, *Pyricularia*, *Ophiostoma*, *Chaetomium*, and *Sordaria* species. Any gene with a hit that was not from *Neurospora* was removed. This analysis results in 670 genes herein termed *Neurospora* LSGs.

Due to the frequent duplication history behind *Neurospora* lineage-specific genes, we enforced a strict expect threshold that identified 670 genes that likely are unique in *Neurospora*. To understand how these 670 LSGs identified from *N. crassa* are shared among the *Neurospora* genomes, a 3-way reciprocal blastp and a tblastx with a PAM30 score matrix were used to search for homologous genes among the three *Neurospora* genomes, and an *E* value of 1 × 10^-10^ was used as a cutoff, and synteny among the orthologs shared within the three *Neurospora* genomes was further visually checked at the FungiDB. Species-specific genes in *N. tetrasperma* and *N. discreta* were not investigated in this study, except for some specific cases mentioned in the text. For uncertain recent duplications, we also relied on the ortholog group identification in FungiDB, which reported potential homologs in 286 fungal and fungus-like Oomycetes genomes, including genomes for 35 Sordariomycetes species closely related to *Neurospora*.

When necessary, additional phylogenetic analyses using the sequences downloaded from the FungalDB homologs groups were pursued to identify lineage-specific genes in *Neurospora*.

### Expression and functional analyses LSGs

Genome-wide gene expression was investigated in *N. crassa* along multiple stages of its life cycle, including conidial germination on different media [23] and production of meiotic propagules (ascospores) on synthetic crossing medium [22]. Transcriptomic data of GSE41484 was reanalyzed with the latest annotation of the *N. crassa* genome. The tally for each sample was processed with LOX v1.6 [94] to analyze gene expression levels across all data points, which uses a Bayesian algorithm to amalgamate different types of datasets. Gene counts or RPKM reads were analyzed by LOX, reporting relative expression of each gene normalized to the lowest treatment, 95% confidence intervals for relative expression, and statistical significance of expression differences. *P* values were adjusted following the procedure of Benjamini et al [99,100]. To assess environmental impacts on expression of LSGs, recent available data on 37 C BM from this lab (GSE168995) and a 240-minute time course of asexual growth in response to darkness and light stimulation [46] were revisited. To assess possible roles of LSGs in metabolic regulation, transcriptomics data from mycelia exposed to 5 different carbon resources from crop residues [39] and from mycelia in response to non-preferred carbon sources such as furfural and 5-hydroxymethyl furfural [HMF; 40] were also examined separately. To assess the gene expression effects of mutations of transcription factors, including *adv-1*, *pp-1*, and *ada-6*, transcriptomics data from Fisher et al. (2018) and Sun et al. (2019) [53,59] were also analyzed with LOX v1.6 [94]. Coordinated expression among genes within the LSG clusters was first manually examined. Twenty-six LSG clusters and their neighbor genes were selected for computation of pairwise correlation coefficients using R. Pairs of genes exhibiting coordination coefficients higher than 0.5 were considered coordinately expressed.

### Clustering analyses of LSGs heterogeneous variation sites and profiling of historical selection

The chromosomal distribution and clustering of LSGs—as well as LSGs and HET-domain genes—were analyzed with Cluster Locator [36]. Cluster Locator requires a parameter (Max-Gap) that specifies the number of genes that can be “skipped” between genes that are considered to be part of a cluster. We set `max-gap = 5`, 1, and 0 (Max-Gap. Statistically significant clusters (*P* < 0.01) were reported. To provide a more continuous measure of gene clustering, a vector of 0s and 1s representing non-LSG genes and LSGs was generated as an input sequence for MACML, a powerful algorithm to profile clustering of discrete ordered data [101]. This algorithm calculates all likely models of linear clustering by partitioning the entire sequence into all possible clusters and subclusters, and all models are statistically evaluated for information-optimality via Akaike Information Criterion [102], ‘corrected’ Akaike Information criterion [103], or Bayesian Information Criterion [104]. In this study, weighted likelihoods were computed based on the conservative Bayesian Information Criterion for each model. LSG cluster probabilities were calculated as weighted averages of models, and ninety-five percent model uncertainty intervals were calculated by further analyses of the model distributions.

To quantify historical selection occurring during divergence of *N. crassa* from its sister species *N. tetrasperma*, window-free regionalized nucleotide site-specific selection profiles were computed on the sequences of LSGs based on the SNP data retrieved from the FungiDB database [105] using Model Averaged Site Selection via Poisson Random Field (MASS-PRF; [106].

MASS-PRF identifies an ensemble of intragenic clustering models for polymorphic (SNPs in *N. crassa*) and divergent sites against orthologs in the *N. tetrasperma* genome (FGSC2508, *mat A* strain, https://mycocosm.jgi.doe.gov/Neute_matA2/Neute_matA2.home) using the Poisson Random Field equations [107] to estimate selection intensity on a regionalized site-by-site basis.

### Knockout strains and phenotype identification

Knockout strains for more than 9600 genes [52], including deletion cassettes for genes in either of the two mating types, were acquired from the Fungal Genetic Stock Center [FGSC: 75]. Identified *Neurospora* LSGs were examined for altered phenotypes during conidia germination on Bird Medium (BM) and sexual development on Synthetic Crossing Medium (SCM) from protoperithecium differentiation to ascospore release. Genotype *mat A* strains were assayed for phenotypes when available; otherwise, *mat a* strains were used. All available KO strains were phenotyped on BM and on SCM with three replicates. For each investigated strain, 3000–5000 conidia were plated onto 90 mm diameter plates and monitored, and crossing was conducted between opposite mating types. Three independent phenotyping experiments were performed with each knockout strain using stored conidia supplied by the FGSC.

## Acknowledgements

We thank the Broad Institute, FungiDB and JGI for making *Neurospora* related fungi genomic data available. All authors have declared that no competing interests exist. This study was supported by funding to JPT from the National Institutes of Health R01 grant AI146584, by the National Science Foundation grant IOS 1457044 to JPT, National Science Foundation IOS 1456482 to FT, and BSF-2018712 to OY. The funders had no role in the study design, data collection and interpretation, or the decision to submit the work for publication.

---

**Figure S1.**
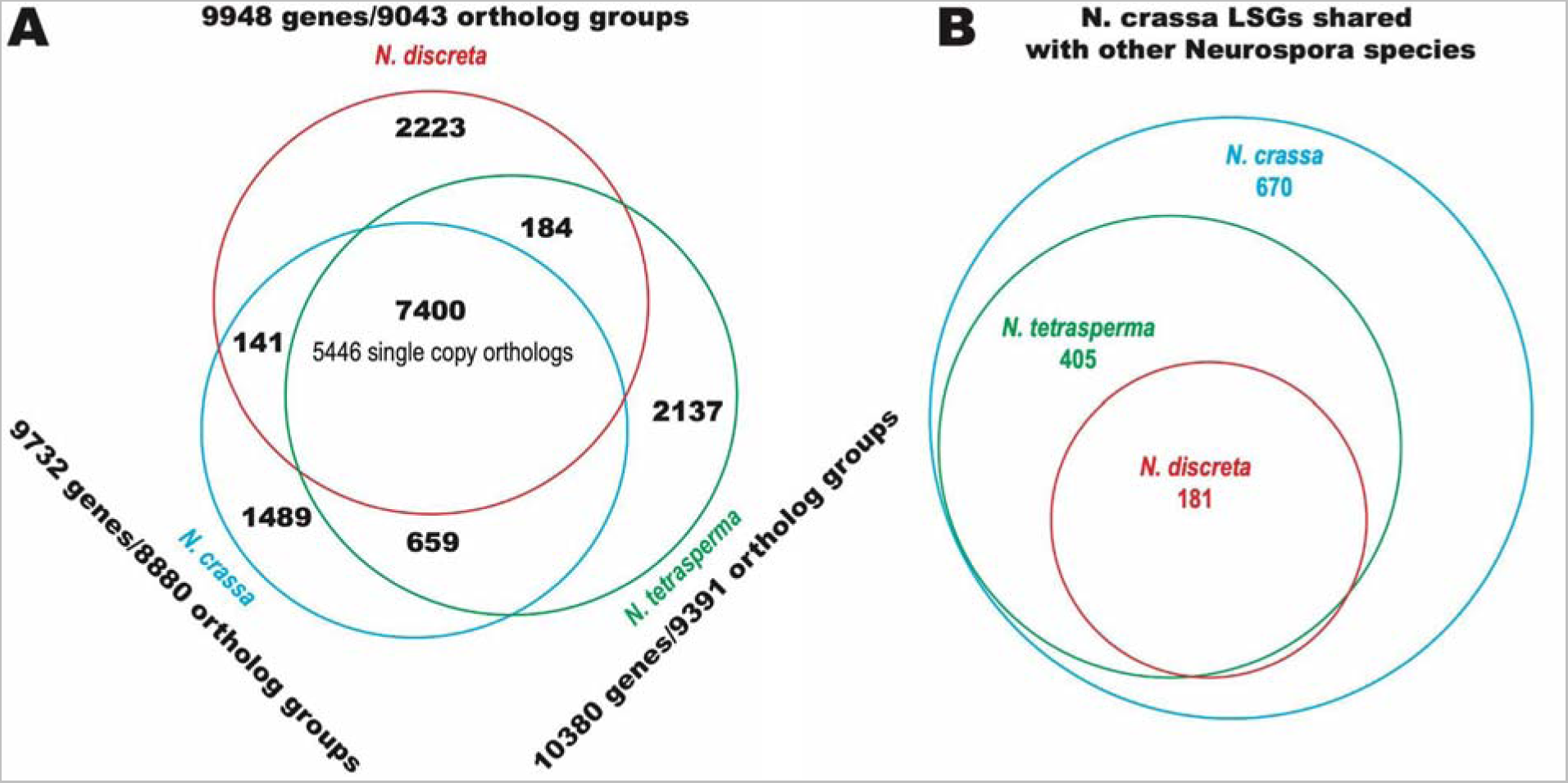
Genome-wide genes and Neurospora LSGs compared within the three Neurospora species N. crassa, N. tetrasperma, and N. discreta. (A) Comparative genomic protein-coding gene content among N. crassa, N. discreta and N. tetrasperma, centering shared single-copy orthologs within the three species. (B) Some Neurospora LSGs in the N. crassa genome are shared within N. tetrasperma and N. discreta genomes.

**Figure S2.**
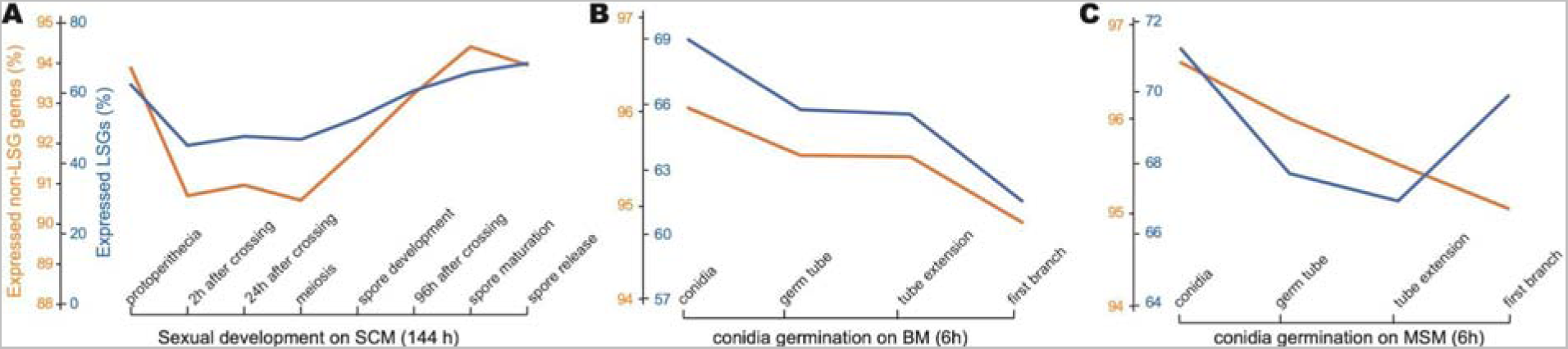
Proportion of *Neurospora* LSGs (blue) and non-LSG genes (orange) that were expressed at sampling points from sexual development on SCM and conidial germination and growth on BM and MSM. (**A**) Sexual development from protoperithecia (starting stage) to mature perithecia at 144 h [22]. (**B**) Asexual growth from conidial germination to the first hyphal branching on Bird medium supporting only asexual development. (C) Asexual growth from conidial germination to the first hyphal branching on maple sap medium supporting both asexual and sexual reproduction [conidial germination; 23].

**Figure S3.**
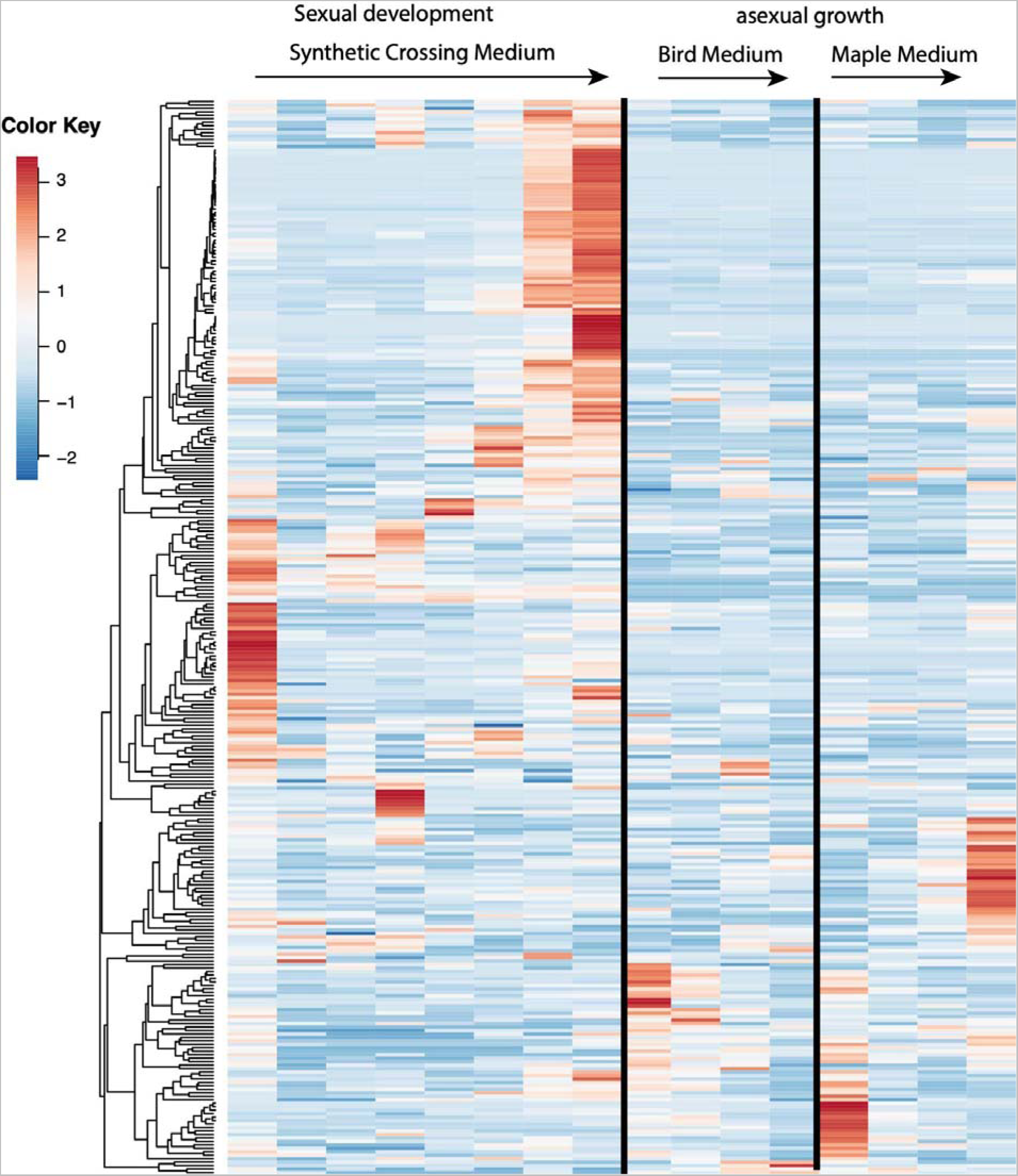
Gene expression dynamics of LSGs across sexual development from protoperithecia (starting stage) to mature perithecia at 144 h [22] and asexual growth from conidial germination to the first hyphal branching, on Bird medium supporting only asexual development, and on a maple sap medium supporting both asexual and sexual reproduction [conidial germination; 23]. Heatmap was generated using the ClustVis web tool.

**Figure S4.**
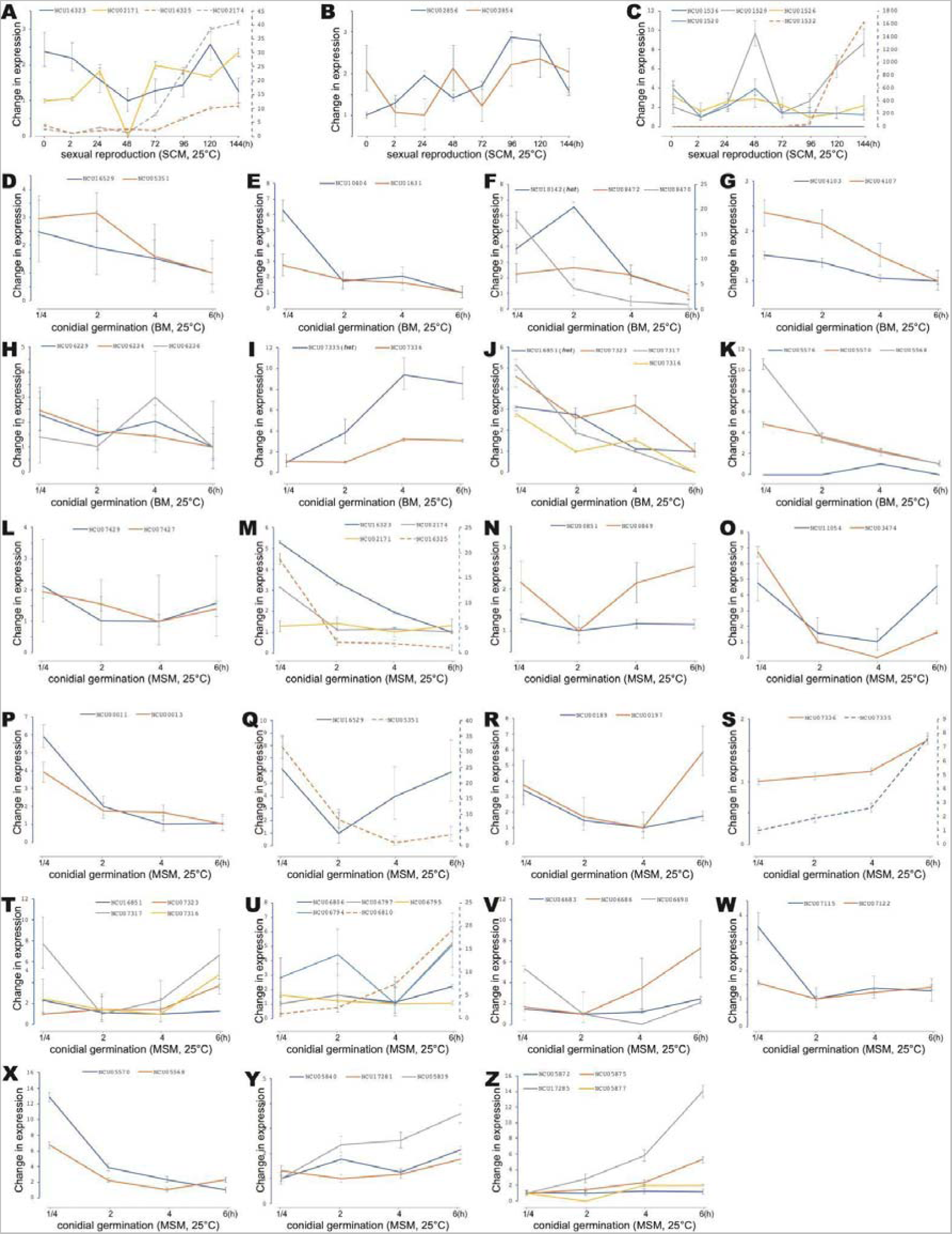
Expression profiles of 21 LSGs clusters and LSG-het gene clusters (Table S4) across asexual and sexual growth in N. crassa. Expression and 95% credible intervals for (A–C) genes in clusters 24, 37, and 131 during sexual development, (D–K) genes in clusters 121, 128, 133, 88, 62, 50, 51 and 8 during conidial germination and asexual growth on Bird medium, and (L–Z) genes in clusters 22, 24, 33, 117, 63, 121, 65, 50, 51, 125, 87, 1, 8, 109 and 110 during conidial germination and asexual growth on maple sap medium.

**Figure S5.**
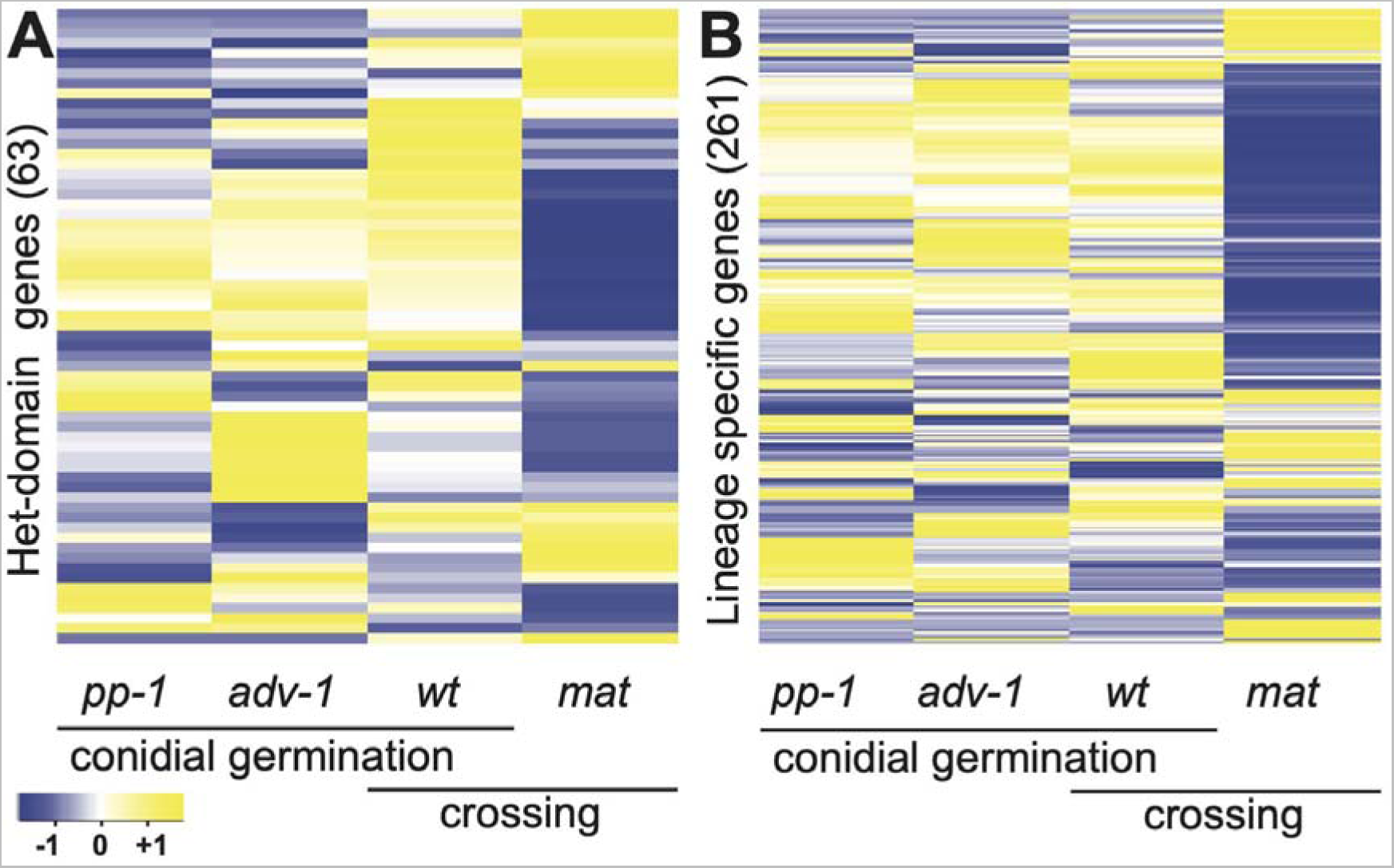
Heatmaps demonstrating divergent expression between wild-type strains and deletions of *pp*-1 and *adv*-1 in their expression of HET-domain and LSGs in conidial germlings, and in mutants of *mat 1-2-1* in fertilized protoperithecia (crossing). (A) Expression divergence of HET-domain genes in three mutants vs. wild type, expression levels sampled in crossing were scaled according to the wild-type expression measured in germlings, and HET-domain genes with 0 measurements in wild type were excluded; (B) Expression divergence of LSGs in three mutants vs. wild type, expression levels sampled in crossing were scaled accordingly to the wild type expression measured in germlings, and LSGs with 0 measurements in wild type were excluded.

**Figure S6.**
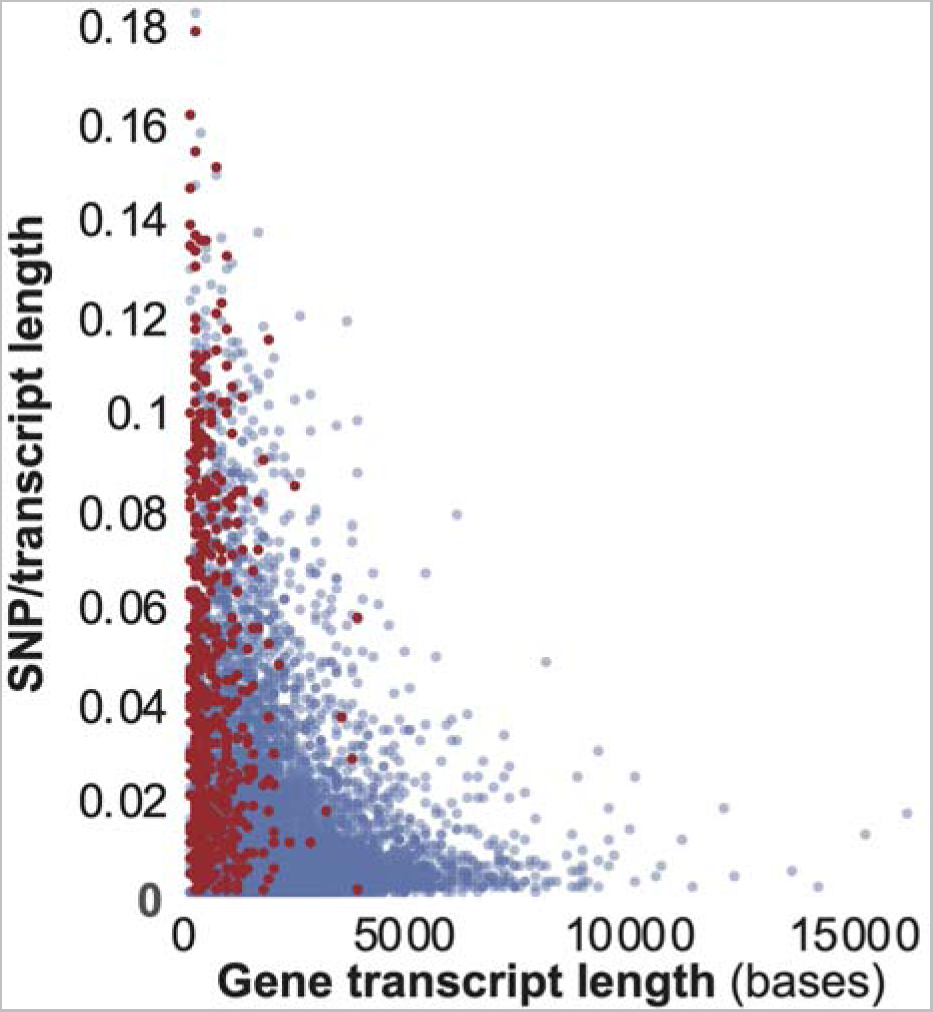
SNPs per nucleotide of transcript—including singletons—for 9756 *N. crassa* genes (LSGs, red; non-LSG, blue) from 26 sequenced strains in FungiDB.

**Figure S7.**
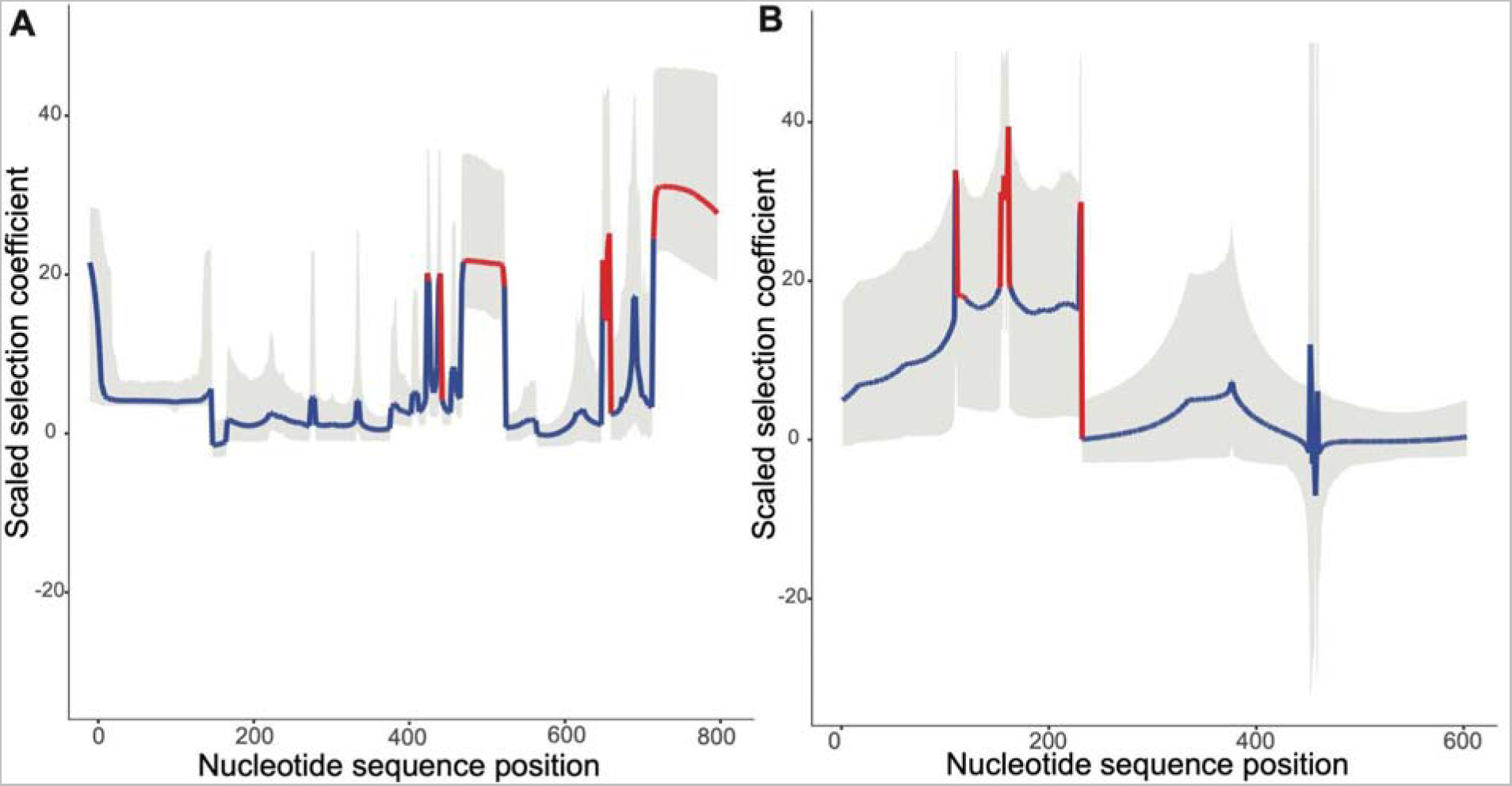
Window-free regionalized nucleotide site-specific selection profiles in two LSGs showing statistically significant gene-regional positive selection along the divergence between *Neurospora crassa* and its sister species *N. tetrasperma*. Scaled selection coefficients (model-averaged *γ*) are plotted (red: lower bound of γ > 4; blue: lower bound of γ ≤ 4; gray band: 95% model uncertainty interval) by nucleotide position for (A) NCU07210, a Neurospora LSG, and (B) NCU02932, a *Neurospora-Sordaria* LSGs.

## References

1. Weisman CM. The Origins and Functions of De Novo Genes: Against All Odds? J Mol Evol. 2022;90: 244–257.

2. Tautz D, Domazet-Lošo T. The evolutionary origin of orphan genes. Nat Rev Genet. 2011;12: 692–702.

3. McLysaght A, Hurst LD. Open questions in the study of de novo genes: what, how and why. Nat Rev Genet. 2016;17: 567–578.

4. Su Z, Townsend JP. Utility of characters evolving at diverse rates of evolution to resolve quartet trees with unequal branch lengths: analytical predictions of long-branch effects. BMC Evol Biol. 2015;15: 86.

5. Dornburg A, Su Z, Townsend JP. Optimal Rates for Phylogenetic Inference and Experimental Design in the Era of Genome-Scale Data Sets. Syst Biol. 2019;68: 145–156.

6. Weisman CM, Murray AW, Eddy SR. Many, but not all, lineage-specific genes can be explained by homology detection failure. PLoS Biol. 2020;18: e3000862.

7. Ruiz-Orera J, Hernandez-Rodriguez J, Chiva C, Sabidó E, Kondova I, Bontrop R, et al. Origins of De Novo Genes in Human and Chimpanzee. PLoS Genet. 2015;11: e1005721.

8. Begun DJ, Lindfors HA, Thompson ME, Holloway AK. Recently evolved genes identified from *Drosophila yakuba* and *D. erecta* accessory gland expressed sequence tags. Genetics. 2006;172: 1675–1681.

9. Begun DJ, Lindfors HA, Kern AD, Jones CD. Evidence for de novo evolution of testis-expressed genes in the *Drosophila yakuba*/*Drosophila erecta* clade. Genetics. 2007;176: 1131–1137.

10. Zhang L, Ren Y, Yang T, Li G, Chen J, Gschwend AR, et al. Rapid evolution of protein diversity by de novo origination in *Oryza*. Nat Ecol Evol. 2019;3: 679–690.

11. Cai JJ, Woo PCY, Lau SKP, Smith DK, Yuen K-Y. Accelerated evolutionary rate may be responsible for the emergence of lineage-specific genes in ascomycota. J Mol Evol. 2006;63: 1–11.

12. Domazet-Loso T, Brajković J, Tautz D. A phylostratigraphy approach to uncover the genomic history of major adaptations in metazoan lineages. Trends Genet. 2007;23: 533– 539.

13. Casola C. From De Novo to “De Nono”: The Majority of Novel Protein-Coding Genes Identified with Phylostratigraphy Are Old Genes or Recent Duplicates. Genome Biol Evol. 2018;10: 2906–2918.

14. Vakirlis N, Carvunis A-R, McLysaght A. Synteny-based analyses indicate that sequence divergence is not the main source of orphan genes. Elife. 2020;9. doi:10.7554/eLife.53500

15. Gladieux P, De Bellis F, Hann-Soden C, Svedberg J, Johannesson H, Taylor JW. Neurospora from Natural Populations: Population Genomics Insights into the Life History of a Model Microbial Eukaryote. In: Dutheil JY, editor. Statistical Population Genomics. New York, NY: Springer US; 2020. pp. 313–336.

16. Davis RH, Perkins DD. Timeline: *Neurospora*: a model of model microbes. Nat Rev Genet. 2002;3: 397–403.

17. Mitchell MB. A model predicting characteristics of genetic maps in *Neurospora crassa*. Nature. 1965;205: 680–682.

18. Ebbole D. Neurospora: a new (?) model system for microbial genetics: Neurospora 2000, Asilomar, CA, USA, 9–12 March 2000. Trends Genet. 2000;16: 291–292.

19. Zhang N, Wang Z. 3 Pezizomycotina: Sordariomycetes and Leotiomycetes. In: McLaughlin DJ, Spatafora JW, editors. Systematics and Evolution: Part B. Berlin, Heidelberg: Springer Berlin Heidelberg; 2015. pp. 57–88.

20. Nowrousian M, Würtz C, Pöggeler S, Kück U. Comparative sequence analysis of *Sordaria macrospora* and Neurospora crassa as a means to improve genome annotation. Fungal Genet Biol. 2004;41: 285–292.

21. Zámocký M, Tafer H, Chovanová K, Lopandic K, Kamlárová A, Obinger C. Genome sequence of the filamentous soil fungus *Chaetomium cochliodes* reveals abundance of genes for heme enzymes from all peroxidase and catalase superfamilies. BMC Genomics. 2016;17: 763.

22. Wang Z, Lopez-Giraldez F, Lehr N, Farré M, Common R, Trail F, et al. Global gene expression and focused knockout analysis reveals genes associated with fungal fruiting body development in *Neurospora crassa*. Eukaryot Cell. 2014;13: 154–169.

23. Wang Z, Miguel-Rojas C, Lopez-Giraldez F, Yarden O, Trail F, Townsend JP. Metabolism and Development during Conidial Germination in Response to a Carbon-Nitrogen-Rich Synthetic or a Natural Source of Nutrition in *Neurospora crassa*. MBio. 2019;10. doi:10.1128/mBio.00192-19

24. Lehr NA, Wang Z, Li N, Hewitt DA, López-Giráldez F, Trail F, et al. Gene expression differences among three *Neurospora* species reveal genes required for sexual reproduction in *Neurospora crassa*. PLoS One. 2014;9: e110398.

25. Wang Z, López-Giráldez F, Wang J, Trail F, Townsend JP. Integrative Activity of Mating Loci, Environmentally Responsive Genes, and Secondary Metabolism Pathways during Sexual Development of *Chaetomium globosum*. MBio. 2019;10. doi:10.1128/mBio.02119-19

26. Corcoran P, Anderson JL, Jacobson DJ, Sun Y, Ni P, Lascoux M, et al. Introgression maintains the genetic integrity of the mating-type determining chromosome of the fungus *Neurospora tetrasperma*. Genome Res. 2016;26: 486–498.

27. Galagan JE, Selker EU. RIP: the evolutionary cost of genome defense. Trends Genet. 2004;20: 417–423.

28. Gladyshev E. Repeat-Induced Point Mutation and Other Genome Defense Mechanisms in Fungi. Microbiol Spectr. 2017;5. doi:10.1128/microbiolspec.FUNK-0042-2017

29. Kasuga T, Mannhaupt G, Glass NL. Relationship between phylogenetic distribution and genomic features in *Neurospora crassa*. PLoS One. 2009;4: e5286.

30. Haridas S, Salamov A, Grigoriev IV. Fungal Genome Annotation. Methods Mol Biol. 2018;1775: 171–184.

31. Grigoriev I. Fungal Genomics Program. Lawrence Berkeley National Lab.(LBNL), Berkeley, CA (United States); 2012. Available: https://www.osti.gov/biblio/1165591

32. Grigoriev IV, Nikitin R, Haridas S, Kuo A, Ohm R, Otillar R, et al. MycoCosm portal: gearing up for 1000 fungal genomes. Nucleic Acids Res. 2014;42: D699–704.

33. Zhao J, Gladieux P, Hutchison E, Bueche J, Hall C, Perraudeau F, et al. Identification of Allorecognition Loci in *Neurospora crassa* by Genomics and Evolutionary Approaches. Mol Biol Evol. 2015;32: 2417–2432.

34. Ament-Velásquez SL, Vogan AA, Granger-Farbos A, Bastiaans E, Martinossi-Allibert I, Saupe SJ, et al. Allorecognition genes drive reproductive isolation in *Podospora anserina*. Nat Ecol Evol. 2022;6: 910–923.

35. Wang Z, Gudibanda A, Ugwuowo U, Trail F, Townsend JP. Using evolutionary genomics, transcriptomics, and systems biology to reveal gene networks underlying fungal development. Fungal Biol Rev. 2018;32: 249–264.

36. Pazos Obregón F, Soto P, Lavín JL, Cortázar AR, Barrio R, Aransay AM, et al. Cluster Locator, online analysis and visualization of gene clustering. Bioinformatics. 2018;34: 3377–3379.

37. Saupe SJ, Kuldau GA, Smith ML, Glass NL. The product of the *het-C* heterokaryon incompatibility gene of *Neurospora crassa* has characteristics of a glycine-rich cell wall protein. Genetics. 1996;143: 1589–1600.

38. Galagan JE, Calvo SE, Borkovich KA, Selker EU, Read ND, Jaffe D, et al. The genome sequence of the filamentous fungus *Neurospora crassa*. Nature. 2003;422: 859–868.

39. Wang B, Cai P, Sun W, Li J, Tian C, Ma Y. A transcriptomic analysis of Neurospora crassa using five major crop residues and the novel role of the sporulation regulator *rca-1* in lignocellulase production. Biotechnol Biofuels. 2015;8: 21.

40. Feldman D, Kowbel DJ, Cohen A, Glass NL, Hadar Y, Yarden O. Identification and manipulation of genes involved in sensitivity to furfural. Biotechnol Biofuels. 2019;12: 210.

41. Eilers FI, Sussman AS. Conversion of furfural to furoic acid and furfuryl alcohol by *Neurospora ascospores*. Planta. 1970;94: 253–264.

42. Emerson MR. Chemical Activation of Ascospore Germination in *Neurospora crassa*. J Bacteriol. 1948;55: 327–330.

43. Kritsky MS, Belozerskaya TA, Sokolovsky VY, Filippovich SY. Photoreceptor Apparatus of the Fungus *Neurospora crassa*. Mol Biol. 2005;39: 514–528.

44. Wang Z, Wang J, Li N, Li J, Trail F, Dunlap JC, et al. Light sensing by opsins and fungal ecology: NOP-1 modulates entry into sexual reproduction in response to environmental cues. Mol Ecol. 2018;27: 216–232.

45. Wang Z, Li N, Li J, Dunlap JC, Trail F, Townsend JP. The Fast-Evolving *phy-2* Gene Modulates Sexual Development in Response to Light in the Model Fungus *Neurospora crassa*. MBio. 2016;7: e02148.

46. Wu C, Yang F, Smith KM, Peterson M, Dekhang R, Zhang Y, et al. Genome-wide characterization of light-regulated genes in *Neurospora crassa*. G3. 2014;4: 1731–1745.

47. Chen C-H, Ringelberg CS, Gross RH, Dunlap JC, Loros JJ. Genome-wide analysis of light-inducible responses reveals hierarchical light signaling in *Neurospora*. EMBO J. 2009;28: 1029–1042.

48. Káldi K, González BH, Brunner M. Transcriptional regulation of the *Neurospora* circadian clock gene *wc-1* affects the phase of circadian output. EMBO Rep. 2006;7: 199–204.

49. Dekhang R, Wu C, Smith KM, Lamb TM, Peterson M, Bredeweg EL, et al. The *Neurospora* Transcription Factor ADV-1 Transduces Light Signals and Temporal Information to Control Rhythmic Expression of Genes Involved in Cell Fusion. G3. 2017;7: 129–142.

50. Fu C, Iyer P, Herkal A, Abdullah J, Stout A, Free SJ. Identification and characterization of genes required for cell-to-cell fusion in *Neurospora crassa*. Eukaryot Cell. 2011;10: 1100– 1109.

51. Lan N, Ye S, Hu C, Chen Z, Huang J, Xue W, et al. Coordinated Regulation of Protoperithecium Development by MAP Kinases MAK-1 and MAK-2 in *Neurospora crassa*. Front Microbiol. 2021;12: 769615.

52. Colot HV, Park G, Turner GE, Ringelberg C, Crew CM, Litvinkova L, et al. A high-throughput gene knockout procedure for *Neurospora* reveals functions for multiple transcription factors. Proc Natl Acad Sci U S A. 2006;103: 10352–10357.

53. Fischer MS, Wu VW, Lee JE, O’Malley RC, Glass NL. Regulation of Cell-to-Cell Communication and Cell Wall Integrity by a Network of MAP Kinase Pathways and Transcription Factors in. Genetics. 2018;209: 489–506.

54. Pöggeler S, Kück U. Comparative analysis of the mating-type loci from *Neurospora crassa* and *Sordaria macrospora*: identification of novel transcribed ORFs. Mol Gen Genet. 2000;263: 292–301.

55. Newmeyer D. A suppressor of the heterokaryon-incompatibility associated with mating type in *Neurospora crassa*. Can J Genet Cytol. 1970;12: 914–926.

56. Jacobson DJ. Control of mating type heterokaryon incompatibility by the *tol* gene in *Neurospora crassa* and *N. tetrasperma*. Genome. 1992;35: 347–353.

57. Xiang Q, Glass NL. The control of mating type heterokaryon incompatibility by vib-1, a locus involved in het-c heterokaryon incompatibility in *Neurospora crassa*. Fungal Genet Biol. 2004;41: 1063–1076.

58. Gearing LJ, Cumming HE, Chapman R, Finkel AM, Woodhouse IB, Luu K, et al. CiiiDER: A tool for predicting and analyzing transcription factor binding sites. PLoS One. 2019;14: e0215495.

59. Sun X, Wang F, Lan N, Liu B, Hu C, Xue W, et al. The Zn(II)2Cys6-Type Transcription Factor ADA-6 Regulates Conidiation, Sexual Development, and Oxidative Stress Response in. Front Microbiol. 2019;10: 750.

60. Ellison CE, Hall C, Kowbel D, Welch J, Brem RB, Glass NL, et al. Population genomics and local adaptation in wild isolates of a model microbial eukaryote. Proc Natl Acad Sci U S A. 2011;108: 2831–2836.

61. Leeder AC, Jonkers W, Li J, Glass NL. Early colony establishment in *Neurospora crassa* requires a MAP kinase regulatory network. Genetics. 2013;195: 883–898.

62. Sun X, Yu L, Lan N, Wei S, Yu Y, Zhang H, et al. Analysis of the role of transcription factor VAD-5 in conidiation of *Neurospora crassa*. Fungal Genet Biol. 2012;49: 379–387.

63. Greenwald CJ, Kasuga T, Glass NL, Shaw BD, Ebbole DJ, Wilkinson HH. Temporal and spatial regulation of gene expression during asexual development of *Neurospora crassa*. Genetics. 2010;186: 1217–1230.

64. Rodriguez S, Ward A, Reckard AT, Shtanko Y, Hull-Crew C, Klocko AD. The genome organization of *Neurospora crassa* at high resolution uncovers principles of fungal chromosome topology. G3. 2022;12. doi:10.1093/g3journal/jkac053

65. Jamieson K, McNaught KJ, Ormsby T, Leggett NA, Honda S, Selker EU. Telomere repeats induce domains of H3K27 methylation in *Neurospora*. Elife. 2018;7. doi:10.7554/eLife.31216

66. Wu C, Kim Y-S, Smith KM, Li W, Hood HM, Staben C, et al. Characterization of chromosome ends in the filamentous fungus *Neurospora crassa*. Genetics. 2009;181: 1129–1145.

67. Mir-Rashed N, Jacobson DJ, Dehghany MR, Micali OC, Smith ML. Molecular and functional analyses of incompatibility genes at het-6 in a population of *Neurospora crassa*. Fungal Genet Biol. 2000;30: 197–205.

68. Xiong Y, Wu VW, Lubbe A, Qin L, Deng S, Kennedy M, et al. A fungal transcription factor essential for starch degradation affects integration of carbon and nitrogen metabolism. PLoS Genet. 2017;13: e1006737.

69. Znameroski EA, Coradetti ST, Roche CM, Tsai JC, Iavarone AT, Cate JHD, et al. Induction of lignocellulose-degrading enzymes in *Neurospora crassa* by cellodextrins. Proc Natl Acad Sci U S A. 2012;109: 6012–6017.

70. Coradetti ST, Craig JP, Xiong Y, Shock T, Tian C, Glass NL. Conserved and essential transcription factors for cellulase gene expression in ascomycete fungi. Proc Natl Acad Sci U S A. 2012;109: 7397–7402.

71. Coradetti ST, Xiong Y, Glass NL. Analysis of a conserved cellulase transcriptional regulator reveals inducer-independent production of cellulolytic enzymes in *Neurospora crassa*. Microbiologyopen. 2013;2: 595–609.

72. Craig JP, Coradetti ST, Starr TL, Glass NL. Direct target network of the *Neurospora crassa* plant cell wall deconstruction regulators CLR-1, CLR-2, and XLR-1. MBio. 2015;6: e01452– 15.

73. Kasuga T, Glass NL. Dissecting colony development of *Neurospora crassa* using mRNA profiling and comparative genomics approaches. Eukaryot Cell. 2008;7: 1549–1564.

74. Bobrowicz P, Pawlak R, Correa A, Bell-Pedersen D, Ebbole DJ. The *Neurospora crassa* pheromone precursor genes are regulated by the mating type locus and the circadian clock. Mol Microbiol. 2002;45: 795–804.

75. McCluskey K, Wiest A, Plamann M. The Fungal Genetics Stock Center: a repository for 50 years of fungal genetics research. J Biosci. 2010;35: 119–126.

76. Herold I, Kowbel D, Delgado-Álvarez DL, Garduño-Rosales M, Mouriño-Pérez RR, Yarden O. Transcriptional profiling and localization of GUL-1, a COT-1 pathway component, in *Neurospora crassa*. Fungal Genet Biol. 2019;126: 1–11.

77. Herold I, Zolti A, Garduño-Rosales M, Wang Z, López-Giráldez F, Mouriño-Pérez RR, et al. The GUL-1 Protein Binds Multiple RNAs Involved in Cell Wall Remodeling and Affects the MAK-1 Pathway in *Neurospora crassa*. Front Fungal Biol. 2021;2. doi:10.3389/ffunb.2021.672696

78. Carrillo AJ, Cabrera IE, Spasojevic MJ, Schacht P, Stajich JE, Borkovich KA. Clustering analysis of large-scale phenotypic data in the model filamentous fungus *Neurospora crassa*. BMC Genomics. 2020;21: 755.

79. Plissonneau C, Stürchler A, Croll D. The Evolution of Orphan Regions in Genomes of a Fungal Pathogen of Wheat. MBio. 2016;7. doi:10.1128/mBio.01231-16

80. de Jonge R, Bolton MD, Kombrink A, van den Berg GCM, Yadeta KA, Thomma BPHJ. Extensive chromosomal reshuffling drives evolution of virulence in an asexual pathogen. Genome Res. 2013;23: 1271–1282.

81. Hartmann FE, Sánchez-Vallet A, McDonald BA, Croll D. A fungal wheat pathogen evolved host specialization by extensive chromosomal rearrangements. ISME J. 2017;11: 1189– 1204.

82. Wacker T, Helmstetter N, Wilson D, Fisher MC, Studholme DJ, Farrer RA. Two-speed genome evolution drives pathogenicity in fungal pathogens of animals. Proc Natl Acad Sci U S A. 2023;120: e2212633120.

83. Shiu PKT, Glass NL. Molecular Characterization of *tol*, a Mediator of Mating-Type-Associated Vegetative Incompatibility in *Neurospora crassa*. Genetics. 1999;151: 545–555.

84. Mitema A, 1 OMICS Research Group, Department of Biotechnology, Vaal University of Technology, Vanderbijlpark, Africa S, et al. Molecular and Vegetative Compatibility Groups Characterization of Aspergillus flavus Isolates from Kenya. AIMS Microbiol. 2020. pp. 231–250. doi:10.3934/microbiol.2020015

85. Grubisha LC, Cotty PJ. Genetic isolation among sympatric vegetative compatibility groups of the aflatoxin-producing fungus *Aspergillus flavus*. Mol Ecol. 2010;19: 269–280.

86. Puhalla JE. Classification of strains of *Fusarium oxysporum* on the basis of vegetative compatibility. Can J Bot. 1985;63: 179–183.

87. Elias KS. Vegetative Compatibility Groups in *Fusarium oxysporum*f. sp.*lycopersici*. Phytopathology. 1991;81: 159.

88. Cai JJ, Petrov DA. Relaxed purifying selection and possibly high rate of adaptation in primate lineage-specific genes. Genome Biol Evol. 2010;2: 393–409.

89. Hartfield M, Poulsen NA, Guldbrandtsen B, Bataillon T. Using singleton densities to detect recent selection in. Evol Lett. 2021;5: 595–606.

90. Ke X, Taylor MS, Cardon LR. Singleton SNPs in the human genome and implications for genome-wide association studies. Eur J Hum Genet. 2008;16: 506–515.

91. Clark TA, Guilmette JM, Renstrom D, Townsend JP. RNA extraction, probe preparation, and competitive hybridization for transcriptional profiling using *Neurospora crassa* long-oligomer DNA microarrays. Fungal Genet Rep. 2008;55: 18–28.

92. Trail F, Wang Z, Stefanko K, Cubba C, Townsend JP. The ancestral levels of transcription and the evolution of sexual phenotypes in filamentous fungi. PLoS Genet. 2017;13: e1006867.

93. Borkovich KA, Alex LA, Yarden O, Freitag M, Turner GE, Read ND, et al. Lessons from the genome sequence of *Neurospora crassa*: tracing the path from genomic blueprint to multicellular organism. Microbiol Mol Biol Rev. 2004;68: 1–108.

94. Zhang Z, López-Giráldez F, Townsend JP. LOX: inferring Level Of eXpression from diverse methods of census sequencing. Bioinformatics. 2010;26: 1918–1919.

95. Arnold R, Rattei T, Tischler P, Truong M-D, Stümpflen V, Mewes W. SIMAP—The similarity matrix of proteins. Bioinformatics. 2005;21: ii42–ii46.

96. Rattei T, Arnold R, Tischler P, Lindner D, Stümpflen V, Mewes HW. SIMAP: the similarity matrix of proteins. Nucleic Acids Res. 2006;34: D252–6.

97. Pellegrini M, Marcotte EM, Thompson MJ, Eisenberg D, Yeates TO. Assigning protein functions by comparative genome analysis: protein phylogenetic profiles. Proc Natl Acad Sci U S A. 1999;96: 4285–4288.

98. Schmid KJ, Aquadro CF. The evolutionary analysis of “orphans” from the *Drosophila* genome identifies rapidly diverging and incorrectly annotated genes. Genetics. 2001. Available: https://www.genetics.org/content/159/2/589.short

99. Benjamini Y, Bretz F, Sarkar SK. Recent Developments in Multiple Comparison Procedures. IMS; 2004.

100. Benjamini Y, Heller R, Yekutieli D. Selective inference in complex research. Philos Trans A Math Phys Eng Sci. 2009;367: 4255–4271.

101. Zhang Z, Townsend JP. Maximum-likelihood model averaging to profile clustering of site types across discrete linear sequences. PLoS Comput Biol. 2009;5: e1000421.

102. H. Akaike. A new look at the statistical model identification. IEEE Trans Automat Contr. 1974;19: 716–723.

103. Hurvich CM, Tsai C-L. Regression and time series model selection in small samples. Biometrika. 1989;76: 297–307.

104. Raftery AE, Madigan D, Hoeting JA. Bayesian Model Averaging for Linear Regression Models. J Am Stat Assoc. 1997;92: 179–191.

105. Stajich JE, Harris T, Brunk BP, Brestelli J, Fischer S, Harb OS, et al. FungiDB: an integrated functional genomics database for fungi. Nucleic Acids Res. 2012;40: D675–81.

106. Zhao Z-M, Campbell MC, Li N, Lee DSW, Zhang Z, Townsend JP. Detection of Regional Variation in Selection Intensity within Protein-Coding Genes Using DNA Sequence Polymorphism and Divergence. Mol Biol Evol. 2017;34: 3006–3022.

107. Sawyer SA, Hartl DL. Population genetics of polymorphism and divergence. Genetics. 1992;132: 1161–1176.

