## Supplementary material for "Lineage-specific genes are clustered with allorecognition loci and respond to G × E factors regulating the switch from asexual to sexual reproduction in *Neurospora*": Tables S13, S14

**Table S13.** Results of MASS-PRF analysis**.** SNPs from 48 strains sampled from natural settings (Ellison et al. 2011), with singletons excluded, were analyzed with MASS-PRF, and some regions of some *Neurospora-Sordaria* orphan genes were inferred to have experienced strong positive or negative selection.

| **Selection intensity** | **Genes ID** |
| --- | --- |
| Strongly positively selected | NCU01032, 01881, 02770, 02932, 05395, 06391, 06759, 06995, 07552, 07618, 07804, 09030, 09562, 09576, 09693, 09824 |
| Moderate positively selected | NCU00496, 01306, 08150, 09137, 00748 |
| Negatively selected | NCU04525, 05145, 08185, 02883, 05135, 00749, 00793, 08449, 04497 |

**Table S14.** Functional categories (GO term) significantly enriched in *Neurospora-Sordaria* orphan genes that were inferred with strong positive selection in Table S12.

| **GO ID** | **GO Term** | **Genes in GO Term** | ***P* value** |
| --- | --- | --- | --- |
| GO:0034599 | response to oxidative stress | NCU02932, 07618, 09562, 09824, | 0.026837 |
| GO:0070887 | response to chemical stimulus | NCU02932, 07618, 09562, 09824, | 0.027488 |
| GO:0050896 | response to stimulus | NCU01881, 02932, 07618, 09562, 09824 | 0.029356 |
| GO:0042221 | response to chemical | NCU02932, 07618, 09562, 09824, | 0.047388 |
